# Dysfunction of microglia-mediated synaptic pruning in Autism spectrum disorder

**DOI:** 10.1101/2025.07.20.665801

**Authors:** Chen Yu, Xiao-Peng Zhang, Wei Wang

## Abstract

Autism spectrum disorder (ASD) has been increasingly associated with abnormalities in synaptic pruning. Although significant progress has been made in elucidating the genetic and immunological underpinnings of ASD, its core pathological mechanisms remain poorly understood. In this study, we developed a network model integrating the inflammatory response pathway in microglia with synaptic signaling pathway to investigate how lipopolysaccharide (LPS) and synaptic activity jointly regulate microglia–mediated synaptic pruning. Our results reveal that pro-inflammatory activation of microglia impairs pruning efficiency under reduced synaptic activity, leading to excessive synaptic accumulation particularly in the prefrontal cortex and contributing to ASD-like phenotypes. Conversely, enhanced synaptic activity partially suppresses LPS-induced synaptic apoptosis and promotes synaptic retention. These findings suggest multiple therapeutic strategies, including targeting pruning-related molecular pathways, mitigating neuroinflammation, and modulating synaptic excitability to alleviate ASD symptoms.

## 1. Introduction

Autism spectrum disorder (ASD) is an abnormal neurodevelopmental condition typically diagnosed in early childhood^1^. According to the 2023 World Health Organization report^2^, approximately 1 in 100 children worldwide is diagnosed with ASD. Patients with ASD often exhibit impairments in social interaction and communication, as well as repetitive and restricted behaviors, all of which pose significant challenges to both the individuals and their families^3,4^. Despite substantial advances in understanding the genetic, neurobiological, and immunological aspects of ASD, its core pathological mechanisms remain unclear. Increasing evidence suggests that individuals with ASD share common molecular abnormalities closely associated with the physiological process of synaptic pruning^5,6^.

Synaptic pruning is a crucial process during neural development that eliminates redundant synaptic connections to optimize neural circuitry^7,8^. Histopathological studies of ASD brain tissue have revealed structural abnormalities, such as cortical thickening and increased dendritic spine density in regions like the prefrontal cortex and temporal cortex^8,9^. Gene expression analyses have also shown upregulation of microglia- and immune-related genes in ASD brains, alongside increased numbers of excitatory neurons^5^. These abnormalities are particularly prominent during key neurodevelopmental windows involving neurogenesis and synaptic pruning^5^. Given the central role of synaptic pruning in shaping functional neural circuits, it is crucial to explore how dysregulation of this process may contribute to the pathogenesis of ASD.

A series of modeling studies have focused on how synaptic pruning shapes neural circuits. For example, differential equation models based on calcium signaling dynamics^10^, spike-timing-dependent plasticity (STDP) models^11,12^, and Hopfield networks have been used to simulate the probabilistic rules governing synapse formation and elimination^13,14^. However, current modeling studies on synaptic pruning are primarily focused at the circuit level, with relatively few addressing the intracellular signaling pathways that govern this process, particularly in the context of synaptic pruning related to ASD pathology.

Synaptic pruning is generally regulated by synaptic activity in conjunction with microglia^15–17^. In terms of synaptic activity, there is long-term depression (LTD), which means that synaptic activity is reduced. In contrast to LTD, long-term potentiation (LTP) implies increased synaptic activity^18^. These two processes can be achieved by relying on stimulus in the form of neurotransmitters, electrical pulses^19^. Furthermore, one of the most consistent findings in mouse models of ASD is dysregulation of LTD, which has been observed in different genetic abnormalities and in different brain regions^19^. Dysregulation of LTD reflects abnormal synaptic pruning, and a shared signaling pathway exists between them^20,21^. Furthermore, recent studies have demonstrated that local activation of the mitochondrial apoptotic pathway can drive the selective pruning of dendrites and dendritic spines via a mechanism dependent on Caspase-3 activity^20^. Moreover, Caspase-3 is required for the elimination of dendritic spines in response to physiologic LTD stimulation^22^. These findings highlight Caspase-3 mediated signaling as a central pathway in synaptic pruning initiated by weaken synaptic activity. Collectively, this evidence underscores the critical role of synaptic activity in regulating structural synaptic remodeling.

Microglia, the brain-resident macrophages, continuously survey their microenvironment and interact with neurons in their resting state^23^. They play a pivotal role in synaptic pruning by phagocytosing superfluous or weakened synapses^24^. Disruption of microglial function can impair the normal progression of synaptic pruning. Tremblay et al. first demonstrated that microglia drive experience-dependent synaptic remodeling^25^, laying the foundation for understanding the regulation of synaptic pruning by microglia. Further research has identified TREM2 (Triggering Receptor Expressed on Myeloid Cells 2), a receptor expressed on microglia, as essential for recognizing and eliminating unnecessary synapses during early brain development^26^.

As observed in *TREM2* knockout mice, loss of TREM2 function disrupts microglia-mediated synaptic refinement and leads to ASD-like behavior^26^. Moreover, after inflammatory stimulus such as Lipopolysaccharide (LPS), microglia transition from a resting state to a pro-inflammatory activated state and release cytokines such as TNFα. This activation can impair their phagocytic capacity and induce neuronal apoptosis^23,27–29^. Collectively, growing evidence indicates that abnormal microglial activity, including dysregulated inflammatory signaling and impaired synaptic pruning, plays a critical role in the pathogenesis of ASD^30–32^. However, the specific roles of neuronal synapses in this process, and how microglia-synapse interactions contribute to aberrant pruning and ASD development, remain important areas for further investigation.

In this study, we used LPS as an external inflammatory stimulus and chemical synaptic stimulus (Synstim) to establish a model of microglia-mediated synaptic pruning integration regulated by synaptic activity in an inflammatory environment and attempted to explain the pathogenesis of ASD in terms of synaptic pruning. The model mainly consists of two modules: LPS-induced microglial proinflammatory activation and regulation of synaptic activity by synaptic stimulus, with an interaction between the two modules. We used ordinary differential equations to describe the network dynamics to explore how these factors affect synaptic pruning. Network parameters were determined by fitting multiple sets of experimental data. Our findings suggest that enhanced excitability of synapses seems to contribute to the progression of ASD by regulating the development of a proinflammatory state of microglia, and also confirm that abnormally pro-inflammatory activated microglia have reduced synaptic pruning capability, leading to delayed pruning of microglia that should be pruned in time, which in turn leads to abnormalities in neural circuits associated with ASD. These findings not only shed light on the synergistic role of inflammation and synaptic activity in microglia function, but also provide new insights into the development of potential therapeutic targets for ASD.

## 2. Model and Methods

### 2.1 Model

Synaptic pruning is initiated by synaptic activity and executed by microglia^7^. Experimental evidence has shown that the absence of TREM2 results in impaired microglia-mediated synaptic pruning^26,31^. Moreover, proinflammatory factors can regulate the expression of TREM2 and promote the development of ASD traits^33^. Based on these findings, we constructed a signal network of microglia-synapse interaction, using the pro-inflammatory factor LPS and synaptic stimulus (Synstim) as external input signals to investigate changes in the microglia-mediated synaptic pruning process.

Within this network, we defined two main modules: one describing the pro-inflammatory activation dynamics of microglia, and the other characterizing the regulation of synaptic activity in response to synaptic stimulus. These modules are interconnected via the PtdSer-TREM2-PSD-95 axis, which represents the execution of synaptic pruning. Postsynaptic density protein-95 (PSD-95) serves as a biomarker to reflect the extent of synaptic pruning, as its levels are positively correlated with the number of synapses^34,35^. Of note, this study focuses primarily on the pruning of excitatory synapses.

#### TREM2-NFκB

TREM2 is a receptor expressed on microglia that modulates immune responses by recognizing various ligands, including lipids, lipoproteins, and nucleic acids^26,36^. For example, the binding of exposed phosphatidylserine (ePtdSer) to TREM2 can initiate a phagocytic program in microglia, a process closely associated with synaptic pruning^37^. In our model, the total amount of TREM2 expressed on microglia is denoted as TREM2_*tot*_, which is dynamically regulated by the internal state of microglia (see Eqs. 5 in the Supplementary Information). Since TREM2 must be activated—typically through ligand binding such as to PtdSer—to initiate downstream signaling, we define TREM2_*ac*_ as the activated form of TREM2 (see Eq. 4 in the Supplementary Information).

In addition to synaptic signals, TREM2 is also modulated by inflammatory stimuli. Acting as an anti-inflammatory factor, TREM2 can inhibit the progression of inflammation. However, upon exposure to pro-inflammatory signals such as LPS, TREM2 expression is subject to antagonistic regulation in microglia.

Specifically, LPS induces the activation of the pro-inflammatory transcription factor NFκB. On one hand, NFκB_*p*_ (activated NFκB) directly suppresses the expression of TREM2; on the other hand, TREM2 negatively regulates NFκB activation by inhibiting its phosphorylation. This reciprocal antagonism forms a positive feedback loop between TREM2 and NFκB signaling^31,38^(See Eqs.2, 4 and 5 in the Supplementary information). NFκB_*p*_ further promotes the transcriptional translation of pro-inflammatory cytokines such as TNFα^39^ (see Eqs.3 in the Supplementary information).

Overall, in our model, microglial pro-inflammatory activation is induced by LPS, which we characterize by measuring the levels of TNFα^40–42^. This part involves two feedback loops: the NFκB-TNFα positive feedback loop and the NFκB-TREM2 positive feedback loop. Here, we consider two forms of NFκB: nonphosphorylated (inactive, NFκB) and phosphorylated (active, NFκB_*p*_), and three forms of TREM2: total TREM2 (TREM2_*tot*_), activated TREM2 (TREM2_*ac*_), and inactive TREM2 (TREM2). Activated NFκB exerts an inhibitory effect on the expression of TREM2, thereby contributing to the disruption of the pro-inflammatory and anti-inflammatory balance within microglia caused by LPS. For convenience, we represent this process as [TREM2_*ac*_] inhibiting NFκB phosphorylation in our model. The effect of NFκB on target gene expression is modeled using a Hill function, with a Hill coefficient of 2, considering NFκB acts as a dimeric transcription factor^43,44^. The total level of NFκB (NFκB_*tot*_) is assumed to be constant and does not change significantly upon LPS stimulus.

#### NFκB-TNFα

LPS induces the expression of tumor necrosis factor-alpha (TNFα) through the activation of NFκB signaling^42^. TNFα subsequently acts in an autocrine manner on microglia to further enhance NFκB activation, thereby forming a positive feedback loop^45^ (see Eqs. 2 and 3 in the Supplementary information). For simplicity, we omitted the intermediate process of TNFα binding to its receptor on the microglial surface, as its exclusion did not significantly affect the model outcomes. Additionally, since other inflammatory cytokines exert effects similar to those of TNFα, their omission is unlikely to influence the overall model behavior. Therefore, to reduce complexity, only TNFα is considered as the representative inflammatory mediator modulating NFκB signaling in the current model.

#### CaMKII_*p*_-PP1

Synaptic chemical stimuli, such as neurotransmitters, are detected by receptors on the synaptic membrane. At excitatory synapses, for instance, NMDA receptors bind glutamate and mediate the influx of Ca^2+^ into the postsynaptic neuron^46^. The incoming Ca^2+^ binds to calmodulin (CaM), which modulates the activity of CaMKII and PP1, thereby regulating synaptic strength^18^. This process is also referred to as the regulation of synaptic plasticity.

Activation of CaMKII and activation PP1_*p*_ can significantly alter synaptic activity^47^. CaMKII is a calmodulin kinase that is activated by calcium ions (Ca^2+^) and calmodulin (CaM) and plays an important role in the nervous system by participating in physiological processes such as synaptic plasticity^47,48^. Inhibition of the CaMKII/CREB axis was found to result in an ASD-like phenotype^49^. PP1 (Phosphatase 1), a dephosphorylase that regulates the activity of proteins by removing phosphate groups from proteins (dephosphorylation), can also be involved in the regulation of synaptic plasticity, and its role in the regulation of synaptic plasticity is opposite to that of CaMKII^50^. Notably, PP1 is the active form, while PP1_*p*_ represents its inactive phosphorylated state. Calcium influx, in combination with CaM, regulates the activation of PP1. Dysregulation of CaMKII and PP1 signaling has been implicated in neurodevelopmental disorders such as ASD^49,51^.

In our model, synaptic stimulus (Synstim) leads to calcium influx through NMDA receptors (NMDAR), activating CaMKII and PP1_*p*_^52^ (see Eqs.10 in the Supplementary information). These proteins are crucial as CaMKII is a key protein kinase and PP1 is a critical phosphatase, both of which regulate synaptic long-term potentiation (LTP) and long-term depression (LTD)^52^. LTD is a process in which synaptic strength decreases over time, and the end result is a decrease in synaptic activity over a long period of time. Meanwhile, LTP is a process in which synaptic strength enhances over time, and the end result is a enhanced synaptic activity over a long period of time. Thus, combining CaMKII and PP1 is able to judge and represent changes in synaptic activity. Due to different calcium-dependent activation thresholds (IC_50_ values), CaMKII and PP1_*p*_ activation timing is not synchronized. Therefore, the value of *J*_*acCack*_ and *J*_*dCapp*_ are different, and the value of the former is set to 4 times the value of the latter (see Table 2 in the Supplementary information).

Here, we consider two forms of CaMKII and PP1_*p*_ respectively: CaMKII (inactive form) and CaMKII_*p*_ (active form), PP1_*p*_ (inactive form) and PP1 (active form)^53^.To be specific, the phosphorylation of CaMKII is regulated by calcium ions, and phosphorylated CaMKII (CaMKII_*p*_) can, in turn, act as an activator of itself, establishing a positive feedback regulatory mechanism^53^. In contrast, the activation state of PP1_*p*_ is also calcium-dependent but is regulated through phosphorylation mediated by CaMKII_*p*_. To capture the dynamic interconversion between the phosphorylated and non-phosphorylated forms of CaMKII and PP1, our model adopts Michaelis–Menten kinetics^53^ (see Eqs. 11 and 12 in the Supplementary Information).

To simplify the model, we omit the intra- and inter-molecular autophosphorylation of CaMKII and intermediate steps in the calcium-induced activation of PP1_*p*_. We also assume that the total concentrations of CaMKII (CaMKII_*tot*_) and PP1 (PP1_*tot*_) remain constant during synaptic stimulation. To simulate the response of CaMKII and PP1_*p*_ to transient calcium signals, we employ Hill functions with coefficients of 4 and 8, respectively, reflecting their high sensitivity and cooperativity to calcium fluctuations^48,53^. Calcium overload is not considered in this model.

#### Caspase-3 activation

Synaptic apoptosis can be triggered by both extrinsic and intrinsic signals^21,54^. Research shows that TNFα^55, 56^and PP1^21,57,58^ promote apoptosis by activating Bax. To simplify the model, we employ a Hill function to represent Bax activation, omitting intermediate signaling steps. Activated Bax facilitates the release of mitochondrial Cytochrome C (CytoC)^59^, which further activates Caspase-3 (Casp3)^59^. Active Casp3 amplifies CytoC release by cleaving its inhibitory factors^60^.

Caspase-3 activity is regulated by X-linked inhibitor of apoptosis protein (XIAP), a member of the inhibitor of apoptosis protein (IAP) family that suppresses apoptosis by directly inhibiting caspases^61^. Studies indicate that CaMKII_*p*_ enhances XIAP stability and expression via the PI3K/AKT signaling pathway^62,63^. To streamline the model, we assume that CaMKII_*p*_ directly promotes XIAP expression, disregarding intermediary components of the PI3K/AKT pathway.

Activated Caspase-3 drives the externalization of PtdSer on the cell membrane. PtdSer is one of the most widely studied synaptic eat-me signals^37,64–66^. Under normal conditions, PtdSer is confined to the cytoplasmic leaflet of the cell membrane by flippase activity. During apoptosis, flippase is inactivated, and scramblase is activated, leading to the exposure of PtdSer on the outer membrane surface^17,22^. Exposed PtdSer (ePtdSer) is recognized by the TREM2 receptor, which is activated into TREM2_*ac*_^67^. Activated TREM2 mediates the phagocytosis of the postsynaptic density protein PSD-95, facilitating synaptic pruning^24^. PSD-95 is a key marker of excitatory synapse abundance, and its reduction reflects active synaptic pruning. In our model, this process is described as an inhibitory effect of TREM2_*ac*_ on PSD-95 levels (see Fig.1).

**Figure 1.**
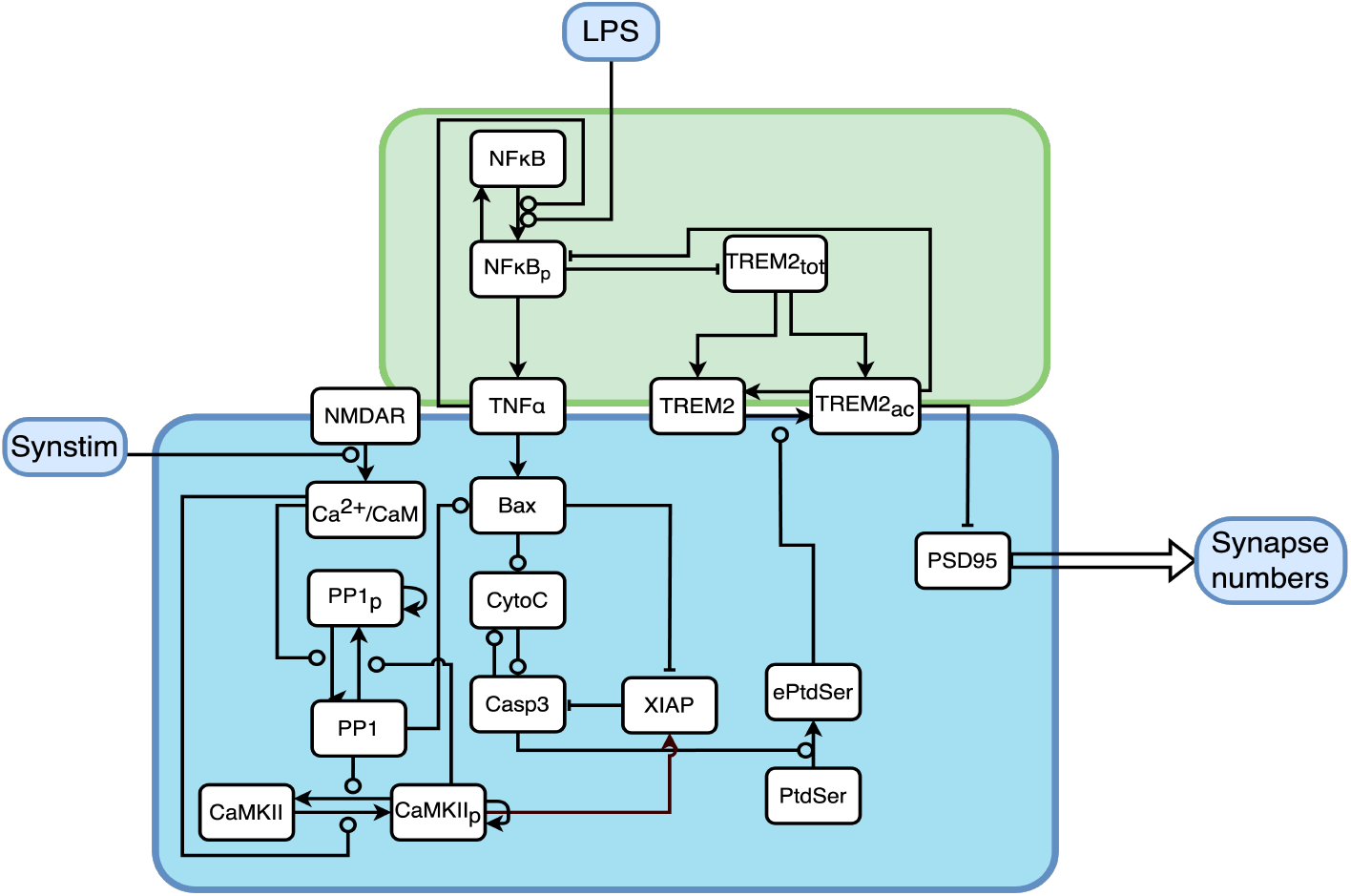
Schematic representation of the model of microglia-mediated synaptic pruning. The model described two parts of the signaling pathways related to synaptic pruning initiation and execution, as well as the pathway module related to the proinflammatory program of microglia. The blue part represents synapses, and the green part represents microglia. On the one hand, LPS induces the phosphorylation of the proinflammation-related transcription factor NFκB in microglia, which promotes the production and release of proinflammatory cytokines TNFα. On the other hand, LPS antagonates the anti-inflammatory program mediated by TREM2, which is highly expressed in resting state, thereby affecting the pro-inflammatory activation of microglia. On the other hand, the synaptic stimulus (Synstim) can modulate the synaptic activity and thus regulate the initiation of the synaptic pruning program. The initiation of apoptosis program will promote the phosphorylation of phosphatidylserine, which binds to TREM2 to perform synaptic pruning. PSD-95 is used as a marker to denote the degree of synaptic pruning.

### 2.1 Method

In the network model, each protein is treated as a state variable, with changes in their concentrations described by ordinary differential equations, detailed in Supplementary Table S1. Phosphorylation and dephosphorylation processes are represented using Michaelis-Menten kinetics, while gene expression and multi-step reaction processes are modeled using Hill functions. The detailed descriptions and initial values of these variables are provided in Supplementary Table S1, and the standard parameters are listed in Supplementary Table S2. The results of sensitivity analysis are also presented in the Supplementary information. Increasing or decreasing the value of the parameter corresponding to 10% has little effect on the movement of the bifurcation point, indicating that the overall sensitivity of the parameter is good (see Supplementary information section 1.3). All protein concentrations are dimensionless. We utilized the free software Oscill8 (available online:http://oscill8.sourceforge.net/) to solve these equations and employed Matlab for visualization.

## 3. Results

### 3.1 Overview-Response of microglia and synapses to LPS

Individuals with ASD display evidence of inflammation, accompanied by increased pro-inflammatory cytokines and microglia activation in the brain^30^. Among these cytokines, Tumor Necrosis Factor-alpha (TNFα) acts as a critical marker of microglial pro-inflammatory activation. By utilizing a branching diagram centered on TNFα, the transitions between different microglial activation states can be effectively visualized, thereby elucidating the neuroinflammatory processes associated with ASD^40^.

As shown in Figure 2A, the steady-state concentration of TNFα ([TNFα]) increases with rising levels of LPS, indicating progressive pro-inflammatory activation of microglia. The steady-state [TNFα] profile delineates four distinct microglial activation states: the resting state (M0), two intermediate or partial pro-inflammatory states (I1-M0/M1 and I2-M0/M1), and the fully activated pro-inflammatory state (M1). These intermediate states are characterized by distinct levels of steady-state [TNFα], reflecting graded activation in response to external stimuli. Variations in synaptic activity are modeled through changes in the parameter Synstim, representing the intensity of synaptic stimulation.

**Figure 2.**
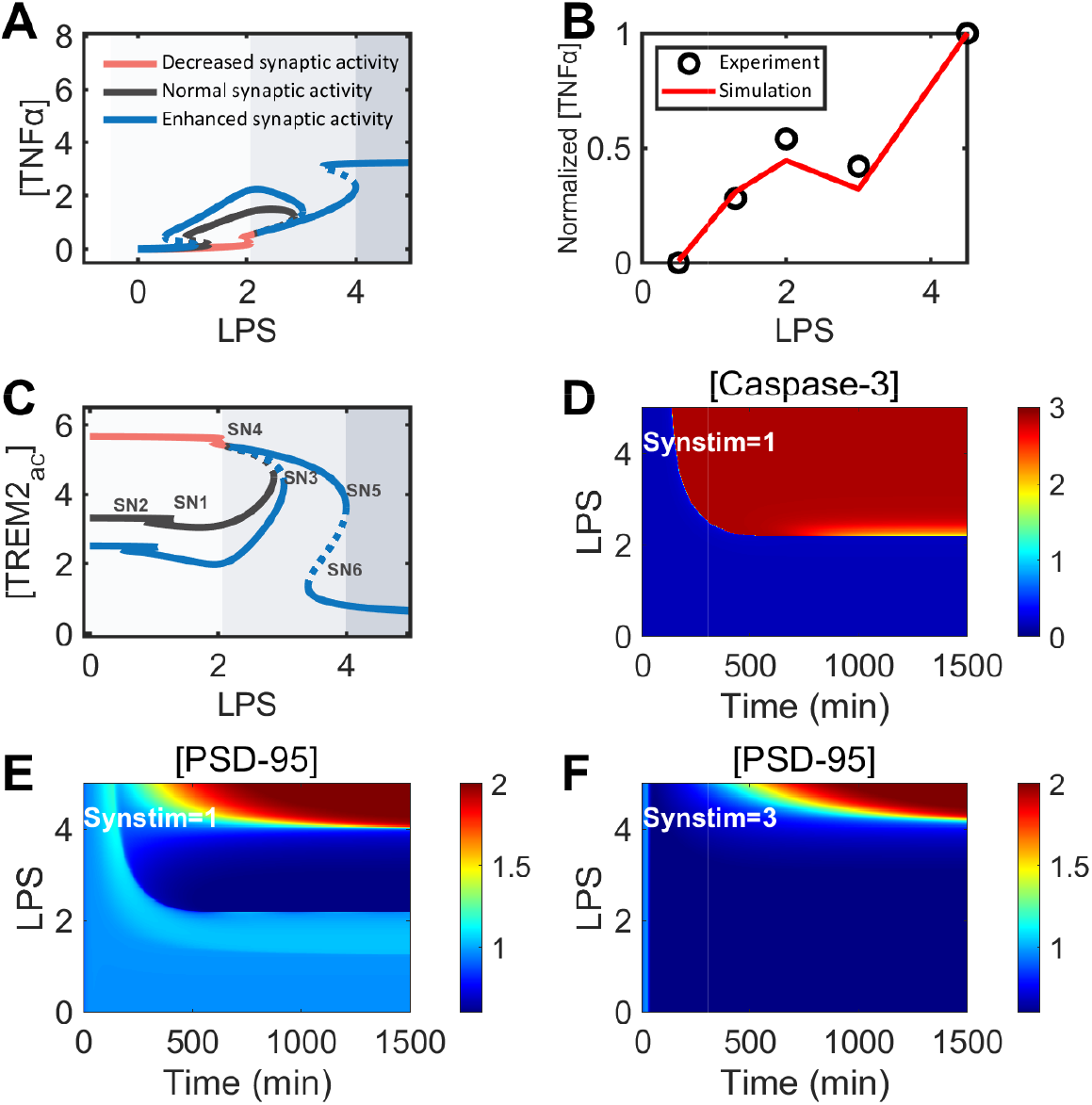
Overview of LPS-regulated changes in microglia and synapses. **A**. Bifurcation analysis of LPS-induced changes in [TNFα] concentration under different synaptic activity conditions. The hollow dots represent saddle node bifurcations. The black line represents the changes under normal synaptic activity (Synstim = 1); the red line represents the changes under weaken synaptic activity (Synstim = 3); the blue line represents the changes under enhanced synaptic activity (Synstim = 5). **B**. Comparison of simulation results and experimental results of LPS-induced changes in [TNFα] concentration under normal synaptic activity. Black represents experimental data, and red represents simulation data^41,42^. (Protein levels are normalized by their respective maximum values). **C**. Bifurcation diagram of LPS-regulated changes in [TREM2_*ac*_] levels under different synaptic activity conditions. The hollow dots are saddle node bifurcations. **D**. Heatmap of [Caspase-3] in response to different degrees of LPS stimulus under normal synaptic activity. **E**. Kinetic heat map of [PSD-95] in response to different degrees of LPS stimulus under normal synaptic activity. **F**. Heatmap of [PSD-95] in response to different degrees of LPS stimulus under weaken synaptic activity.

Figure 2B presents the normalized comparison between simulated outcomes and experimental data. In this figure, the points represent experimental observations, while the lines correspond to model simulations. As the concentration of LPS increases, the steady-state [TNFα] initially rises, then decreases, and subsequently rises again. This non-monotonic pattern is broadly consistent with experimental findings^41,42^. Notably, unlike the monotonic increase in [TNFα] observed when microglia are stimulated in isolation, this non-linear behavior may be attributed to the modulatory influence of neurons. The underlying mechanisms responsible for this interaction warrant further investigation.

Under inflammatory stimulation, such as exposure to LPS, microglia not only initiate pro-inflammatory responses but also activate anti-inflammatory programs, particularly through the TREM2-dependent signaling pathway^38^. This pathway serves to counteract the pro-inflammatory effects of LPS. Concurrently, activation of TREM2 signaling promotes microglia-mediated synaptic pruning. Due to its dual roles in modulating inflammation and regulating synaptic pruning, TREM2 serves as a critical mediator in the pathophysiology of ASD. Therefore, in our model, the activated form of TREM2 (TREM2_*ac*_) is employed as an indicator to reflect both anti-inflammatory responses and synaptic pruning capability.

As illustrated in Figure 2C, under conditions of normal synaptic activity (indicated by the black line), the concentration of TREM2_*ac*_ ([TREM2_*ac*_]) exhibits a non-monotonic response to increasing concentrations of LPS—initially rising and subsequently declining. This trend highlights the complex regulation of TREM2 under inflammatory conditions and suggests that microglia-mediated synaptic pruning reaches its peak at a moderate LPS concentration. Beyond this threshold, further increases in LPS stimulation lead to a significant decline in pruning capability. Furthermore, under conditions of heightened synaptic activity, synaptic pruning is inhibited at low concentrations of LPS, suggesting that increased synaptic input can attenuate microglial pruning capability. This observation aligns with findings in individuals with ASD, where pro-inflammatory microglial activation coincides with an elevated density of dendritic spines, indicating a disruption in the equilibrium between synaptic elimination and maintenance.

As shown in Figures 2A and 2C, after the synaptic activity is enhanced, the first pair of saddle node bifurcation points (SN1-SN2) shift to the left (the saddle node bifurcation points are consistent with those in Figure 2C), indicating that lower LPS stimulus can promote the transition of microglia from the resting state (M0) to the incomplete pro-inflammatory activation state 1 (I1-M0/M1). At the same time, in microglia at this stage, the [TREM2_*ac*_] level decreases with the increase of synaptic activity, reflecting the decrease of synaptic pruning capability, and further suggesting that the increase of synaptic activity can inhibit the pruning capability of resting microglia. In other words, after the synaptic activity is enhanced, the degree of synaptic pruning by microglia is reduced. However, it is worth noting that after continuing to increase the LPS stimulus, the synaptic pruning capability of microglia under conditions of enhanced synaptic activity and normal synaptic activity gradually tends to be consistent, and also experiences a process of first increasing and then decreasing. This similar pruning pattern, on the one hand, shows that the regulation of synaptic activity on microglia is limited and cannot completely offset the effect of LPS on the change of microglial state; on the other hand, it shows that high levels of LPS stimulus will inhibit the pruning capability of microglia.

Similar to normal synaptic activity (Synstim = 1) and enhanced synaptic activity (Synstim = 5), after synaptic activity is weakened, the pruning capability of microglia is also reduced under high-level LPS stimulus, and the three curves converge and overlap after the LPS stimulus level exceeds 3 (dimensionless), once again emphasizing the inhibitory effect of medium and high levels of LPS stimulus on the synaptic pruning capability of microglia. However, unlike normal synaptic activity and enhanced synaptic activity, after synaptic activity is weakened, under the stimulus of low concentrations of LPS, the [TREM2_*ac*_] level increases significantly, and the pruning capability of microglia is significantly enhanced, which may only be related to the activity of the synapse itself. Because under extremely low concentrations of LPS stimulus (LPS=0-1), microglia are in a resting state, and the microglia at this time do not undergo pro-inflammatory activation, which corresponds to microglia in a physiological state. In other words, microglia at this stage can perform synaptic pruning normally. Moreover, when synaptic activity is weakened, it will promote the pruning level of microglia.

In the experiment, LPS stimulus of microglia can also lead to synaptic apoptosis and changes in the number of synapses in neurons co-cultured with microglia. Among them, Caspase-3 (Casp-3) activation can be used as a marker of synaptic apoptosis, while PSD-95 can characterize the number level of excitatory synapses. Therefore, we further explored the direct effects of LPS on [Caspase-3] and [PSD-95] (see Figure2(D-F)). Figure 2D shows the dynamic changes of synapses [Caspase-3] over time under normal synaptic activity conditions. After treatment with medium and high LPS (approximately LPS>2.2), the dynamic changes of [Caspase-3] in synapses changed from low state to high state, indicating that a certain concentration of LPS can induce synaptic apoptosis, and as the LPS concentration gradually increased (LPS>2.2), the time required for [Caspase-3] to reach the high state gradually shortened.

In addition, we compared [PSD-95] levels under normal and weaken synaptic activity. Simulation results under normal synaptic activity (Figure 2E) showed that [PSD-95] exhibited a biphasic, nonlinear response to increasing LPS concentrations. When the LPS concentration ranged from 0 to 2.2 (dimensionless units), [PSD-95] remained stable at approximately 1.0. As LPS increased from 2.2 to 4, [PSD-95] gradually declined over time, reaching a steady-state value of 0.1. However, when LPS exceeded 4, [PSD-95] increased markedly, with the steady-state level rising to 2.0. This biphasic patterninitial decrease followed by a sharp increase–highlights the non-monotonic and complex regulation of synaptic density by LPS, and suggests that microglia in distinct activation states exert different degrees of pruning on excitatory synapses, as reflected by changes in [PSD-95]. Under weaken synaptic activity, the [PSD-95] response to LPS changes markedly (Figure 2F). Across the LPS range of 0–4, [PSD-95] remains consistently low (about 0.1). However, when LPS exceeds 4, [PSD-95] rapidly increases to a high level. Compared to the normal synaptic activity condition, weakened synaptic activity significantly delays the increase in [PSD-95] in response to high LPS concentrations (LPS > 4). This suggests that full pro-inflammatory activation of microglia under conditions of reduced synaptic activity promotes synaptic accumulation, which is consistent with the observed increase in excitatory synapses, such as dendritic spines, in individuals with ASD.

In summary, this series of research results show that high-level LPS can induce pro-inflammatory activation of microglia and inhibit the pruning capability of microglia, which will eventually lead to the emergence of the key pathology of ASD—an abnormal increase in the number of synapses. However, the specific mechanism by which LPS regulates microglia-mediated synaptic pruning is still not fully understood, and the intermediate process of inducing ASD is also unclear. For example, why does the pruning capability of microglia show nonlinear changes under different concentrations of LPS? How does synaptic activity shape the response pattern of microglia to LPS?

### 3.2 LPS modulates microglial inflammation and synaptic Pruning

Experiments have shown that LPS influences both microglia and synapses^41,42^, and can reduce total TREM2 levels ([TREM2_*tot*_]), a finding that is also replicated in our simulations. However, the mechanism by which LPS promotes microglial pro-inflammatory activation and subsequently modulates pruning capability remains unclear. Experimental evidence suggests that inhibition of NFκB activation increases the expression of [TREM2_*tot*_]^43^, while activation of TREM2 inhibits the proinflammatory response of microglia. This suggests that there is a feedback loop of mutual inhibition between NFκB and TREM2. Based on this, we focused on the interplay between the NFκB-centered pro-inflammatory pathway and the TREM2-mediated anti-inflammatory pathway.

When LPS = 5, we simulated the dynamics of [TREM2_*tot*_] by varying the parameter *k*_*acLPSn*_, which represents the LPS-dependent NFκB phosphorylation rate (Figure 3A). The simulation results, under normalized conditions, were consistent with experimental data^43^. Throughout this analysis, synaptic activity was assumed to be at normal levels. Under the baseline setting (*k*_*acLPSn*_ = 0.04), microglia underwent progressive pro-inflammatory activation, accompanied by a reduction in [TREM2_*tot*_], indicating a decrease in their pruning capability (Figure 3A, top). When *k*_*acLPSn*_was reduced to 0.03, the steady-state level of [TREM2_*tot*_] increased, suggesting that inhibition of NFκB phosphorylation could preserve TREM2 expression and thereby restore microglial pruning capability (Figure 3A, bottom). To further explore this relationship, we constructed bifurcation diagrams of [TREM2_*tot*_] and [NFκB] as functions of *k*_*acLPSn*_(Figure 3B). As *k*_*acLPSn*_ increased, [TREM2_*tot*_] decreased, confirming an inverse relationship between NFκB_*p*_ and TREM2 expression. These findings suggest that modulating NFκB phosphorylation may partially reverse the impairment in microglial pruning caused by pro-inflammatory activation.

**Figure 3.**
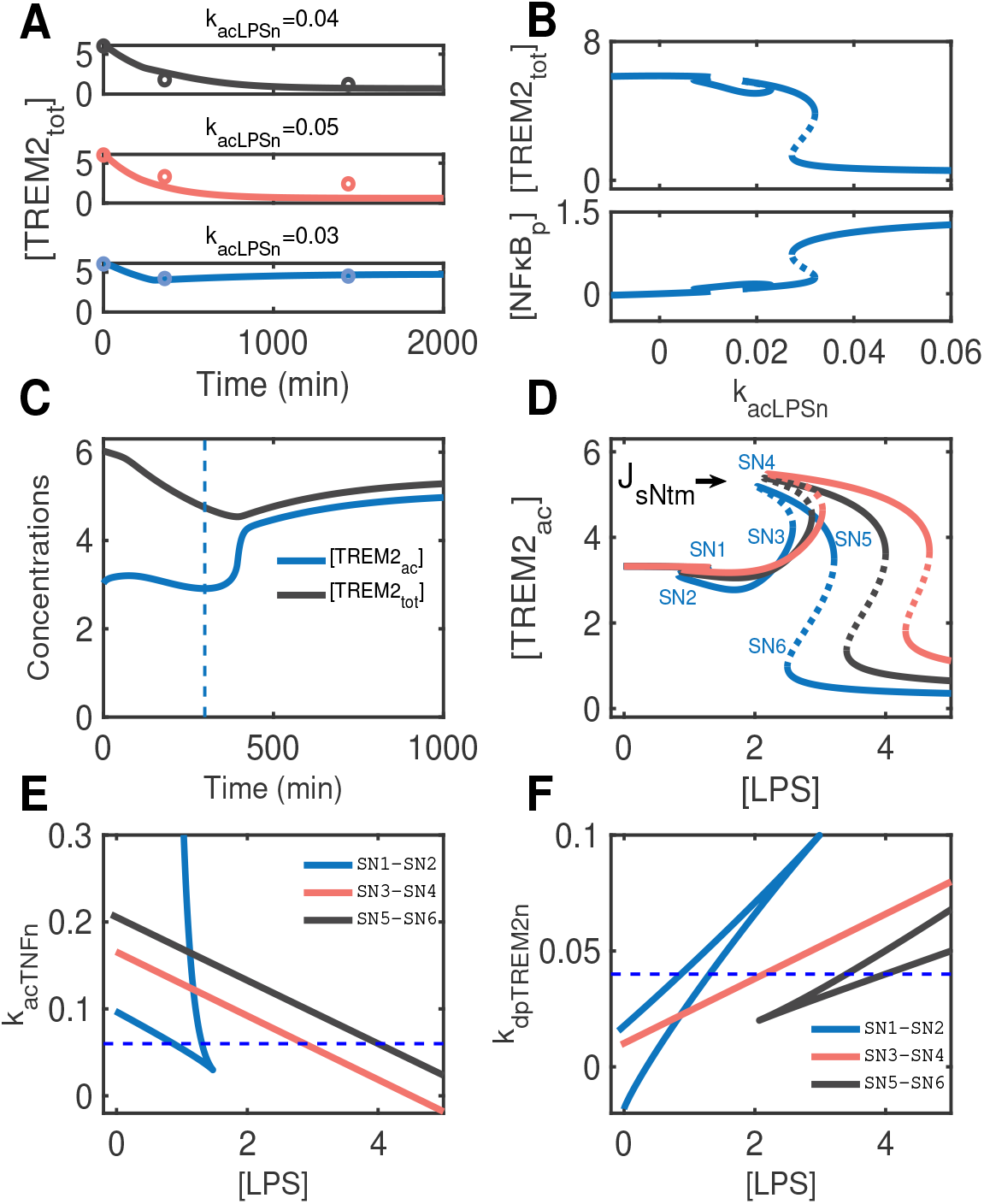
Under normal synaptic activity, LPS contributes to the pro-inflammatory activation of microglia and modulates their synaptic pruning capacity. **A**. Time-course dynamics of [TREM2_*tot*_] at varying values of *k*_*acLPSn*_ (0.03, 0.04 [standard], and 0.05). **B**. Bifurcation diagram of [TREM2_*tot*_] as a function of *k*_*acLPSn*_ under high LPS conditions (LPS = 5). **C**. Temporal dynamics of [TREM2_*ac*_] and [TREM2_*tot*_] with LPS = 3 under normal synaptic activity. **D**. Bifurcation diagram of [TNFα] versus LPS, showing the effect of varying *J*_*sNtm*_ under normal synaptic activity. **E**. Two-parameter analysis of LPS and *k*_*acTNFn*_. **F**. Two-parameter analysis of LPS and *k*_*dpTREM*2*n*_.

When LPS = 3, the dynamics of [TREM2_*ac*_] and [TREM2_*tot*_] are shown in Figure 3C. The dotted lines indicate the time point at which [TREM2_*ac*_] begins to rise. Following this increase, [TREM2_*tot*_] gradually elevates, suggesting that TREM2 activation precedes and potentially promotes its own expression. This implies that TREM2 protein levels can be indirectly upregulated by activation. Furthermore, activation of TREM2 appeared to exert similar effects as inhibition of NFκB activation, both contributing to the restoration of microglia pruning capacity (Figure 3C). These results indicate that, in the absence of full pro-inflammatory activation, enhancing TREM2 activation alone is sufficient to recover microglial function, akin to suppressing NFκB signaling. In other words, early anti-inflammatory intervention, either through the use of TREM2 agonists or by targeting NFκB phosphorylation, may prevent complete pro-inflammatory activation of microglia and thus reduce the risk of ASD.

However, it should be noted that there are also significant differences between these two treatments, and this process directly acting on TREM2 activation may lead to excessive microglia pruning capability compared to inhibiting NFκB phosphorylation. For example, under the normal synaptic activity conditions mentioned above, when LPS = 3, the level of [PSD-95] is significantly lower than when LPS = 1, that is, the synapse is overpruned (see Figure 2E), which is consistent with observations in the brains of individuals with ASD. In other words, improperly administered TREM2 agonists may lead to overpruning of synapses at early developmental stages, resulting in phenotypes of excessive synaptic loss in ASD or Schizophrenia.

Subsequently, to further explore the role of the inhibitory strength of NFκB on TREM2_*tot*_ expression in regulating microglia pruning capability under different LPS stimulus, We performed the simulations by tuning the value of the parameter *J*_*sNtm*_ (the Michaelis constant produced by the NFκB-dependent inhibition of TREM2_*tot*_). As shown in Figure 3D, as *J*_*sNtm*_ increases (from 0.3 to 0.4 to 0.5, the standard value is 0.4), SN3-SN4 and SN5-SN6 move to the right, requiring higher concentrations of LPS to reduce TREM2 and greatly reduce microglia pruning capability. In other words, clinically, increasing the inhibitory threshold of NFκB on TREM2 expression can delay the full proinflammatory activation of LPS dependent microglia, improve the pruning ability of microglia in the process of ASD, and thus alleviate the symptoms of ASD.

Given the network topology (Fig.1) and LPS-induced changes in TREM2 and TNFα levels, NFκB may play a critical role in proinflammatory activation of microglia. Also, two positive feedback loops associated with NFκB may be involved in the formation of bifurcation points. However, how different positive feedback loops affect each bifurcation point remains unclear.

To explore the reason for the bifurcation point formation, we performed a two-parameter analysis with *k*_*acTNFn*_ (TNFα-dependent NFκB activation rate) and LPS as horizontal and vertical coordinates, respectively (Fig.3E). It can be seen from the figure that under the standard parameter value setting, the curve corresponding to SN1-SN2 is non-monotonic and has two intersection points with the dotted line (*k*_*acTNFn*_=0.06), indicating that [TNFα] is involved in the formation of the bifurcation point of this pair. In contrast, SN3-SN4 and SN5-SN6 are monotonically varying and have only one intersection with the dashed line, respectively. However, this monotonous variation also indicates that [TNFα] can affect the movement of bifurcation points, although it does not affect the formation of bifurcation points. It follows that [TNFα] determines the transition of microglial proinflammatory activation from the M0 state to the I1-M0/M1 state and significantly affects the process of microglia from the I2-M0/M1 state to full proinflammatory activation.

Next, we further explore the effect of TREM2 on bifurcation points. Since TREM2 is highly expressed in resting microglia (M0 microglia), it can resist the activation of inflammation related transcription factors such as NFκB^43^. We continue to explore the role of TREM2 in bifurcation point formation by varying the value of the parameter *k*_*dpTREM*2*n*_. *k*_*dpTREM*2*n*_ is fixed to 0.04 (the standard value of the model, see the dotted line in Figure 3F). According to figure 3F, *k*_*dpTREM*2*n*_ has two intersections with SN1-SN2 and SN5-SN6, respectively. This indicates that TREM2 is involved in the formation of these two pairs of bifurcation points. In contrast, SN3-SN4 formation was not affected by TREM2. These results suggest that TREM2 can regulate microglial proinflammatory activation and mainly regulate its transition from M0 to I1-M0/M1 state and from I2-M0/M1 to M1 state. Moreover, SN1-SN2 and SN3-SN4 were found to be sensitive to NFκB activation and TREM2 expression inhibition, as well as to synaptic apoptosis-related parameters (see section 1.3 in supplementary information). SN5-SN6 were mainly sensitive to the parameters related to the positive feedback between TREM2 and NFκB, indicating that the change of microglia pruning ability from high to low was mainly related to the transition of microglia to full proinflammatory activation (see section 1.3 in supplementary information).

In summary, [TNFα] is mainly involved in forming the first pair of bifurcation points, and [TREM2] is mainly involved in forming the third pair of bifurcation points. However, neither of these is directly responsible for the formation of the second pair of bifurcation points, and the formation of SN3-SN4 needs to be further explored.

### 3.3 Regulation of synaptic activity by synaptic stimulus (Synstim)

Synaptic activity is generally modulated in response to external stimuli. The form of synaptic stimulation can be either pulsed electrical stimulation or chemical stimulation. However, any form of synaptic stimulation affects the activity and stability of the synapse itself by changing the hierarchical transmission of signaling pathways within the synapse^68^. A key regulatory module within the synaptic signaling network is the CaMKII–PP1_*p*_ feedback loop, which plays a central role in maintaining synaptic activity and stability. Synaptic stimulus regulates this positive feedback loop by modulating intracellular Ca^2+^ concentrations, leading to long-lasting alterations in synaptic function^47^. These changes in synaptic activity form the molecular basis of microglia-mediated synaptic pruning, suggesting that the relative concentrations of active CaMKII and phosphorylated PP1 (PP1_*p*_) are closely associated with synaptic elimination processes^47^.

Moreover, increasing evidence links dysregulation of CaMKII-PP1_*p*_ dependent synaptic activity to the pathophysiology of ASD. For example, mutations in CaMKII-encoding genes, including *CAMK2A* and *CAMK2B*, have been implicated in increased the probability of ASD^69^. CaMKII is abundantly localized in the postsynaptic density and is crucial for the phosphorylation of AMPA receptors and other synaptic proteins, thereby modulating synaptic strength. Similarly, variants in genes encoding regulatory subunits of PP1 have also been associated with elevated ASD susceptibility^70^.

Synaptic pruning has been shown to share key signaling pathways with long-term depression (LTD), a process characterized by the sustained weakening of synaptic strength^7^. LTD is generally associated with decreased synaptic activity, and both theoretical and experimental studies have demonstrated that intracellular Ca^2+^ concentrations critically determine the induction of either LTP or LTD corresponding to synaptic strengthening or weakening, respectively^71^.

CaMKII and PP1 are both subject to autophosphorylation and engage in a mutually inhibitory feedback loop, which, as captured in our model (Fig. 1), gives rise to tristability in synaptic states. This dynamic allows the synapse to reside in one of three stable activity states, depending on the balance between these key regulators. To simulate synaptic stimulus, we introduced the variable Synstim, which represents the magnitude of external synaptic input. Synstim corresponds to chemical stimulus such as experimentally applied neurotransmitters, which modulates intracellular Ca^2+^ levels via activation of NMDA receptors. Initially, Synstim was set to a minimal value to reflect a basal, physiologically quiescent synaptic state^48^. In our model, Synstim and Ca^2+^ levels are linearly coupled, such that increasing Synstim results in proportionally elevated intracellular Ca^2+^ concentrations. It is important to note that the current model does not account for calcium overload or excitotoxicity induced by elevated intracellular calcium levels.

As shown in Fig.4A and Fig.4B, under negligible synaptic stimulus, both [CaMKII_*p*_] and [PP1] remain at low levels, representing the basal synaptic state characterized by normal physiological activity. Under moderate levels of Synstim, [PP1] transitions to a high state while [CaMKII_*p*_] remains low, indicative of weakened synaptic activity, consistent with LTD-like signaling. If this state becomes dysregulated, it is likely to impair synaptic pruning and thereby contribute to the development of ASD. In contrast, high Synstim levels drive [CaMKII_*p*_] into a high state and suppress [PP1] to a low state, corresponding to enhanced synaptic activity. The simulated data in Fig. 4A–B are in qualitative agreement with previous experimental observations^48^, supporting the models capacity to capture key features of synaptic signaling dynamics across different stimulus regimes.

**Figure 4.**
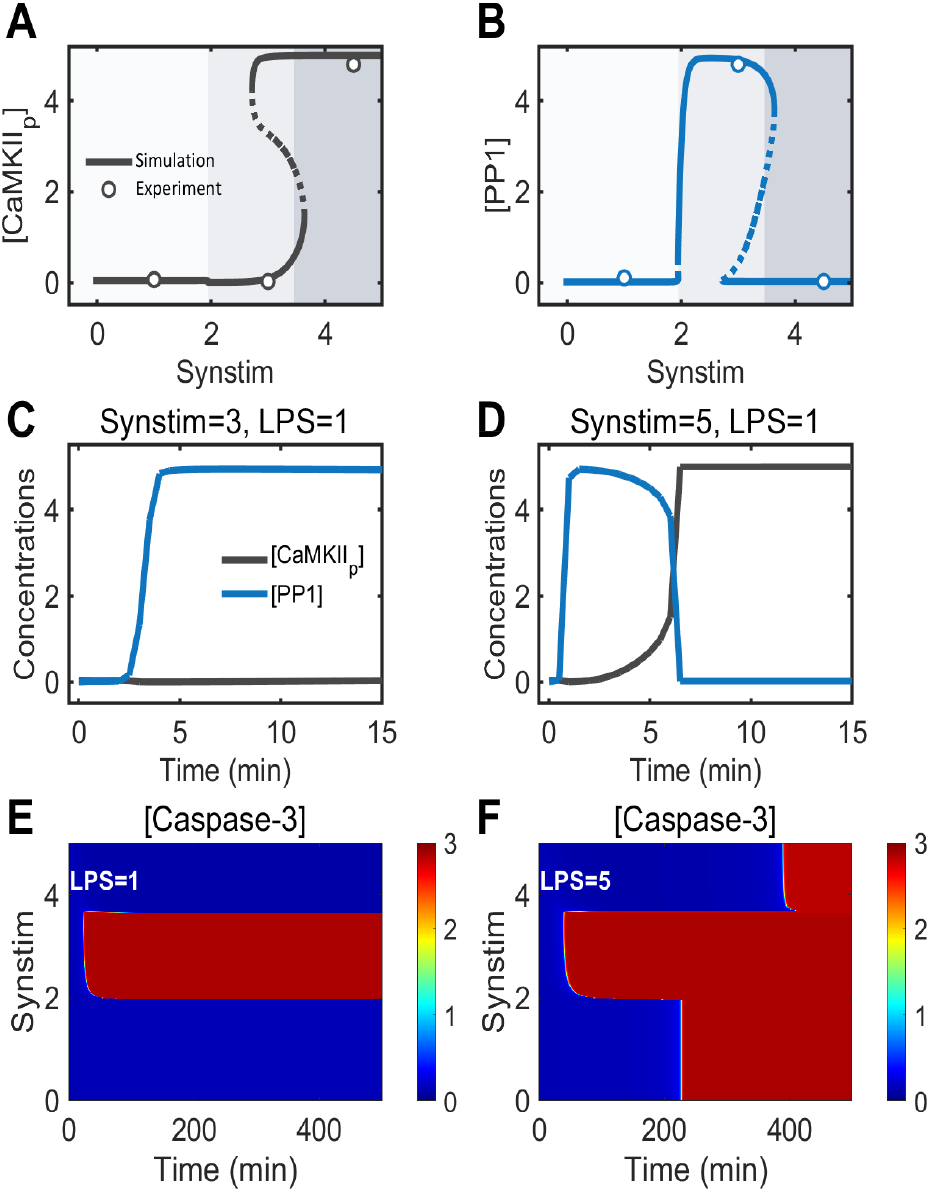
Synaptic stimulus (Synstim) modulates synaptic activity. **A**. Bifurcation diagram of [CaMKII_*p*_] versus Synstim at LPS = 0.1; black dots represent experimental data. **B**. Bifurcation diagram of [PP1] versus Synstim at LPS = 0.1; blue and black dots represent experimental data^48^. **C**. Time course of [CaMKII_*p*_] and [PP1] dynamics at Synstim = 3. **D**. Time course of [CaMKII_*p*_] and [PP1] dynamics at Synstim = 5. **E**. Temporal dynamics of [PSD-95] under varying Synstim levels at low LPS stimulus (LPS = 1). **F**. Temporal dynamics of [PSD-95] under varying Synstim levels at high LPS stimulus (LPS = 5). Note: At LPS = 0.1 and LPS = 1, microglia is considered to be in a resting state.

Next, we sought to further investigate how the aforementioned feedback loop influences downstream synaptic Caspase-3 and the potential role of LPS in this process (Figure 4(E-F)). Figure 4E illustrates the temporal changes in Caspase-3 activity under a low LPS concentration (LPS = 1). When the synaptic stimulus intensity (Synstim) ranges between 2 and 4, Caspase-3 activity transitions from a low to a high state over time. However, when Synstim falls below 2 or exceeds 4, Caspase-3 activity remains in a low state. These findings suggest that, under low LPS conditions, only moderate-intensity synaptic stimulus (Synstim) effectively triggers Caspase-3 activation, whereas both low-and high-intensity stimuluss fail to do so.

Moreover, in addition to applying moderate-intensity Synstim, elevating LPS concentrations can also induce Caspase-3 activation under both low and high Synstim conditions, ultimately reaching a high state. Notably, under low Synstim conditions, the activation of Caspase-3 occurs more rapidly, whereas under high Synstim conditions, this activation is markedly delayed. This phenomenon suggests two key insights: first, the activation of Caspase-3 may directly stem from the accumulation of TNFα resulting from pro-inflammatory microglial activation (as depicted in Figure 1, where TNFα released by microglia acts on synapses); second, enhanced synaptic activity may delay synaptic apoptosis, a finding consistent with established experimental perspectives.

### 3.4. Effects of synaptic Caspase-3 activation on microglial pruning capability

In Section 3.2, we posited that the second pair of bifurcation points observed in the TREM2 diagram is unlikely to be solely driven by LPS-induced microglial regulation, but is more plausibly modulated by synaptic activity. Therefore, in this section, we examine in detail how synaptic activity contributes to the emergence of this bifurcation pair, specifically through Caspase-3-mediated regulation of microglial pruning capability.

As shown in Fig.5A and Fig.5B, when Synstim = 1, [Caspase-3] transitions from a low to a high state as LPS concentration increases, indicating the onset of synaptic apoptosis. In contrast, under Synstim = 3, [Caspase-3] remains in a high state even at low LPS levels, suggesting that the synapse is already in a pro-pruning state.

**Figure 5.**
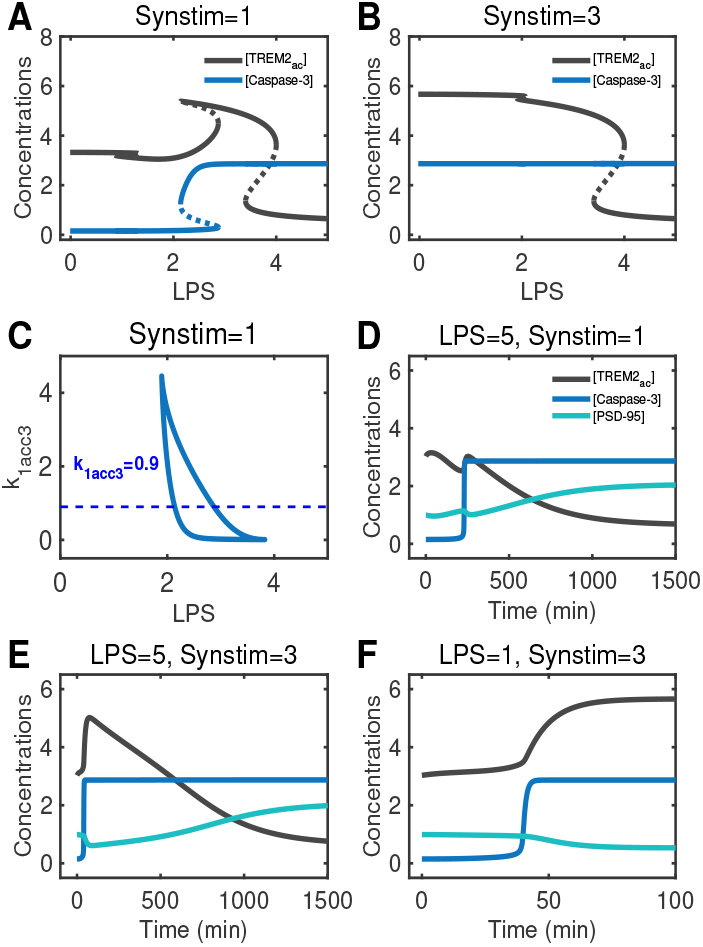
Effects of synaptic Caspase-3 activation on microglial pruning capability. **A**. When Synstim = 1 (normal synaptic activity), the bifurcation diagram of [Caspase-3] (blue) and [TREM2_*ac*_] (black) with LPS. **B**. When Synstim = 3 (weakened synaptic activity), the bifurcation diagram of [Caspase-3] (blue) and [TREM2_*ac*_] (black) with LPS. **C**. When Synstim = 1 (normal synaptic activity), the two-parameter bifurcation diagram of *k*_*acc*3_ and LPS at the second pair of bifurcation points. **D**. When Synstim = 1, LPS = 5 (synaptic activity is normal and microglia are in a fully pro-inflammatory state), the evolution of [TREM2_*ac*_], [Caspase-3], and [PSD-95] over time. **E**. When Synstim = 3, LPS = 5 (synaptic activity is weakened and microglia are in a fully pro-inflammatory state), the evolution of [TREM2_*ac*_], [Caspase-3], and [PSD-95] over time. **F**. When Synstim = 3, LPS = 1 (synaptic activity is weakened and microglia are in a resting state), the evolution of [TREM2_*ac*_], [Caspase-3], and [PSD-95] over time.

Correspondingly, under Synstim = 1, microglial [TREM2_*ac*_] displays a non-monotonic response to LPS. Specifically, within the range of LPS = 0–1.2, microglia exhibit normal pruning capability; from LPS = 1.2–2.8, pruning capability initially decreases and then increases; and in the range of LPS = 2.8–4, although pruning capability gradually declines with further increases in LPS, it remains above the baseline level. Under Synstim = 3, [TREM2_*ac*_] is markedly elevated within the LPS = 0–1.2 range, likely due to Caspase-3 activation triggered by weaken synaptic activity.

To further investigate the link between the second bifurcation pair (SN3–SN4) and Caspase-3 activation, we examined the effect of varying the Caspase-3 activation coefficient, *k*_*acc*3_. As shown in Fig. 5C, when *k*_*acc*3_ = 0.9, the curve intersects the response surface at two points, indicating that Caspase-3 contributes to the elevation of [TREM2_*ac*_] levels. This suggests that during the intermediate state of pro-inflammatory microglial activation, Caspase-3 activation can facilitate an increase in microglial pruning capability.

Following our initial exploration of how LPS and synaptic activity modulate microglial pruning via Caspase-3, we further investigated the time-dependent dynamics of synapse quantity under conditions of normal versus weakened synaptic activity. As shown in Fig. 5D (LPS = 5, Synstim = 1), we examined the temporal changes in [TREM2_*ac*_], [PSD-95], and [Caspase-3] concentrations following full pro-inflammatory activation of microglia under physiologically normal synaptic activity.

At LPS = 5, Caspase-3 activation was observed in both normally active and hypoactive synapses (Fig.5E). However, the final [PSD-95] levels were elevated compared to those under low LPS stimulus (LPS = 1) (Fig.5F). For normally active synapses, Caspase-3 activation may occur in a spatially restricted manner, potentially triggering compensatory mechanisms that counteract synaptic pruning pressure. In such cases, synapses may increase [PSD-95] expression as a protective response to maintain structural and functional integrity.

In contrast, hypoactive synapses, which are already primed for apoptosis, undergo further degeneration upon LPS = 5. Although these synapses reach a pro-apoptotic state, the concomitant high LPS levels downregulate microglial pruning capability, preventing efficient removal of dying synapses. This mismatch leads to pathological accumulation of [PSD-95], reflecting the failure to eliminate dysfunctional synapses. Moreover, compared to normal synaptic activity, hypoactive synapses exhibited consistently higher [TREM2_*ac*_] levels over time, suggesting that weaken synaptic activity may demand a stronger microglial pruning response (compare black curves in Fig.5D and Fig.5E).

In summary, moderate levels of Synstim regulate synaptic pruning by regulating Caspase-3-dependent synaptic apoptosis.

### 3.5. Microglial responses to varying synaptic stimuli under partial pro-inflammatory activation

The emergence of the second pair of bifurcation points is a key determinant of the differential synaptic pruning capability observed in microglia under partial pro-inflammatory activation, characterized by the presence of two intermediate states. In the previous section, we demonstrated that the formation of this second bifurcation is closely associated with Caspase-3 activation and analyzed how synapse numbers evolve under normal versus weaken synaptic activity. However, the functional consequences of this bifurcation on microglial behavior have yet to be fully elucidated.

Moreover, as shown in Fig. 2A, identical steady-state levels of [TNFα] can correspond to distinct microglial states when Synstim is set to either 1 or 5. These states exhibit different pruning capacities, suggesting the involvement of regulatory mechanisms that are not yet fully understood. In this section, we systematically investigate the responses of synapses and microglia under varying Synstim levels, focusing on partially activated microglia. Our aim is to characterize the dynamic behaviors of these distinct intermediate states, identify functional differences between them, and explore how such disparities may contribute to neurodevelopmental disorders such as ASD.

To dissect the differences between the two intermediate microglial states, we examined the temporal dynamics of key molecular components within the system (Fig.6). Figures 6A–6E illustrate the time-course responses of various species over a 1500-minute simulation period under LPS = 3 and three synaptic activity conditions (Synstim = 1, 3, and 5). Each curve represents the temporal evolution of a particular component, enabling comparative analysis across different microglial states. Notably, under weaken synaptic activity (Synstim = 3), the system exhibits only one partially pro-inflammatory microglial state across the LPS range, as previously shown in Fig.2. In Fig.6A, Caspase-3 levels rapidly increase and stabilize at a high plateau in response to Synstim = 3. In contrast, for Synstim = 1 and Synstim = 5, Caspase-3 activation is delayed. These results suggest that Caspase-3 activation is highly sensitive to changes in synaptic input and plays a central role in mediating downstream responses. Furthermore, in conditions of normal or elevated synaptic activity (Synstim = 1 or 5), the delayed Caspase-3 activation is primarily triggered by the microglial transition from the resting state to the second partially pro-inflammatory intermediate state. Consistent with these findings, Fig.6B shows that [TREM2_*ac*_] exhibits a transient peak under Synstim = 3, likely driven by the early burst of Caspase-3 activity, before gradually returning toward baseline. This surge in [TREM2_*ac*_] in turn induces a sharp decrease in [PSD-95], as shown in Fig.6C (blue curve). In contrast, under Synstim = 1 or 5, [TREM2_*ac*_] first decreases and then increases, eventually reaching the same steady-state level observed in the Synstim = 3 condition. These results indicate that, despite the reduced synaptic input, microglia in the partially activated intermediate state 2 can maintain appropriate pruning capability, thereby preserving synaptic homeostasis.

**Figure 6.**
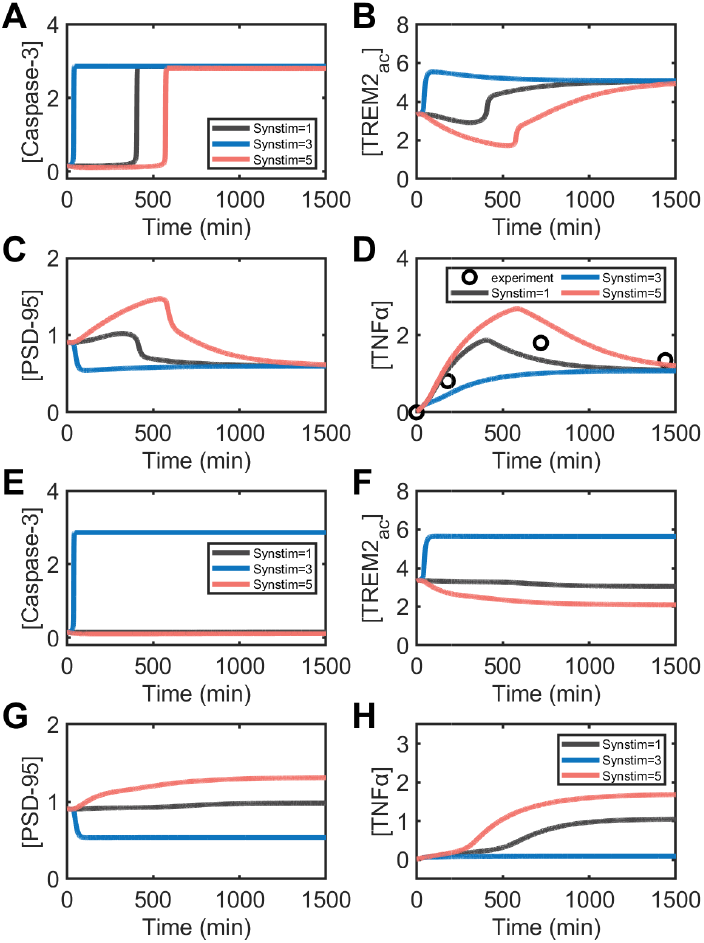
Microglial responses to varying synaptic stimuli under partial pro-inflammatory activation. **A**. Time-course of [Caspase-3] levels under different synaptic stimuli (Synstim) at LPS = 3. **B**. Temporal dynamics of [TREM2_*ac*_] under varying Synstim at LPS = 3. **C**. Temporal dynamics of [PSD-95] under varying Synstim at LPS = 3. **D**. Temporal dynamics of [TNFα] under varying Synstim at LPS = 3. **E**. Time-course of [Caspase-3] levels under varying Synstim at LPS = 1.55. **F**. Temporal dynamics of [TREM2_*ac*_] under varying Synstim at LPS = 1.55. **G**. Temporal dynamics of [PSD-95] under varying Synstim at LPS = 1.55. **H**. Temporal dynamics of [TNFα] under varying Synstim at LPS = 1.55.

The elevated steady-state level of [TREM2_*ac*_] observed at 1500 minutes reflects ongoing synaptic apoptosis. Microglia in incomplete pro-inflammatory activation state 2 (I1-M0/M2) remain responsive to apoptotic cues and are capable of initiating pruning programs, ultimately resulting in a reduction of synaptic density.

Figure 6D depicts the temporal dynamics of [TNFα] under different synaptic activity conditions, showing good agreement with experimental data from Lund et al.^72^ And similar phenomena have also been observed in patients with ASD^73^. The dynamic profile of [TNFα] reflects the precise temporal course of microglial pro-inflammatory activation. On the one hand, [TNFα] levels increase with elevated LPS concentrations, but exhibit a subsequent decline coinciding with Caspase-3 activation. This adaptive response suggests a feedback mechanism whereby Caspase-3 positively regulates microglial activation by constraining the continued rise of [TNFα]. On the other hand, the [TNFα] peak occurs earlier and reaches a lower maximum under Synstim = 1 compared to Synstim = 5, indicating that increased synaptic activity may enhance microglial pro-inflammatory responses. In other words, while elevated synaptic activity attenuates synaptic pruning, it concurrently promotes pro-inflammatory microglial activationa dual effect that may contribute to microglia-mediated inflammation and excitatory synapse accumulation observed in ASD.

Compared to microglia in incomplete pro-inflammatory activation state 2 (I1-M0/M2), those in state 1 exhibit distinct behavior under Synstim = 1 (normal activity) or Synstim = 5 (enhanced activity). In these cases, synaptic Caspase-3 is not activated, suggesting that no negative feedback loop is formed between microglia and synapses. Therefore, although the steady-state level of [TNFα] under Synstim = 1 is similar between the two microglial states, the underlying regulatory mechanisms differ significantly.

To further investigate the mechanistic differences across microglial activation states, we plotted the temporal dynamics of key components under LPS = 1.55 (Figure 6E–6H). Notably, under Synstim = 3, Caspase-3 is activated due to weaken synaptic activity, leading to an elevated level of [TREM2_*ac*_]. In this condition, the synaptic pruning capability of microglia closely resembles that of microglia in the resting state (Figure 6B). Under Synstim = 1, [TREM2_*ac*_] exhibits minimal changes; however, when Synstim = 5, [TREM2_*ac*_] shows a marked decline, accompanied by a noticeable increase in [PSD-95] levels compared to the normal activity condition. These results indicate that, when microglia are in the incomplete pro-inflammatory activation state 1 (I1-M0/M1), enhanced synaptic activity promotes the accumulation of synapses while simultaneously exacerbating the degree of microglial pro-inflammatory activation. This observation may reflect a potential pathological mechanism in ASD, whereby elevated synaptic activity contributes to disease progression. Moreover, it lends further support to the excitatory/inhibitory imbalance hypothesis of ASD, in this case highlighting the contribution of excessive excitation.

In addition, based on the combination of Figure 2A and Figure 6H, microglia remain in a resting state under LPS = 1.55. This indirectly suggests that synaptic pruning induced by weaken synaptic activity may inhibit the pro-inflammatory activation of microglia to some extent, further emphasizing the beneficial regulatory role of microglia-mediated synaptic pruning in dampening neuroinflammation. Specifically, the initiation of early synaptic pruning may trigger rapid NFκB activation while suppressing TNFα production in microglia.

Taken together, these findings highlight the complex and dynamic interactions between microglia and neuronal synapses. When microglia reside in incomplete pro-inflammatory activation state 1 (I1-M0/M1), weaken synaptic activity appears to delay pro-inflammatory activation, while enhanced synaptic activity facilitates inflammation and reduces pruning efficiencypotentially contributing to microglial hyperactivation and excitatory synapse accumulation in ASD. In contrast, when microglia transition to incomplete pro-inflammatory activation state 2 (I2-M0/M1), changes in synaptic activity (whether enhanced or reduced) have diminished regulatory effects and fail to prevent either inflammatory progression or increased pruning capability. These results underscore the critical importance of identifying and targeting “therapeutic windows” for effective intervention in ASD.

### 3.6. LPS and Synstim co-regulate microglial pruning

As can be seen from the previous, microglia with different parts of proinflammatory activation state differ in their synaptic pruning capability, and the initiation of pruning is influenced by synaptic activity. We then further examined how LPS and Synstim co-regulate microglia pruning in ASD.

Tian et al.^31^ reported that [PSD-95] levels are elevated in ASD, that TREM2 overexpression reduces [PSD-95], and that TREM2 silencing further increases it. To compare our model with these experimental findings, we varied the parameters *k*_*acPStm*_ (the *PtdSer*-dependent rate of TREM2 activation) and *k*_*sNtm*_ (the NFκB-dependent inhibition rate of TREM2_*tot*_ synthesis). All other data were normalized using the control group as a reference. As shown in Fig. 7A, [PSD-95] was significantly elevated in the ASD group relative to controls, consistent with experimental results^31^ and pathological observations. TREM2 overexpression in the ASD model reduced [PSD-95] to near-normal levels, whereas TREM2 knockdown led to a marked increase in [PSD-95], underscoring the essential role of TREM2 in synaptic pruning.

**Figure 7.**
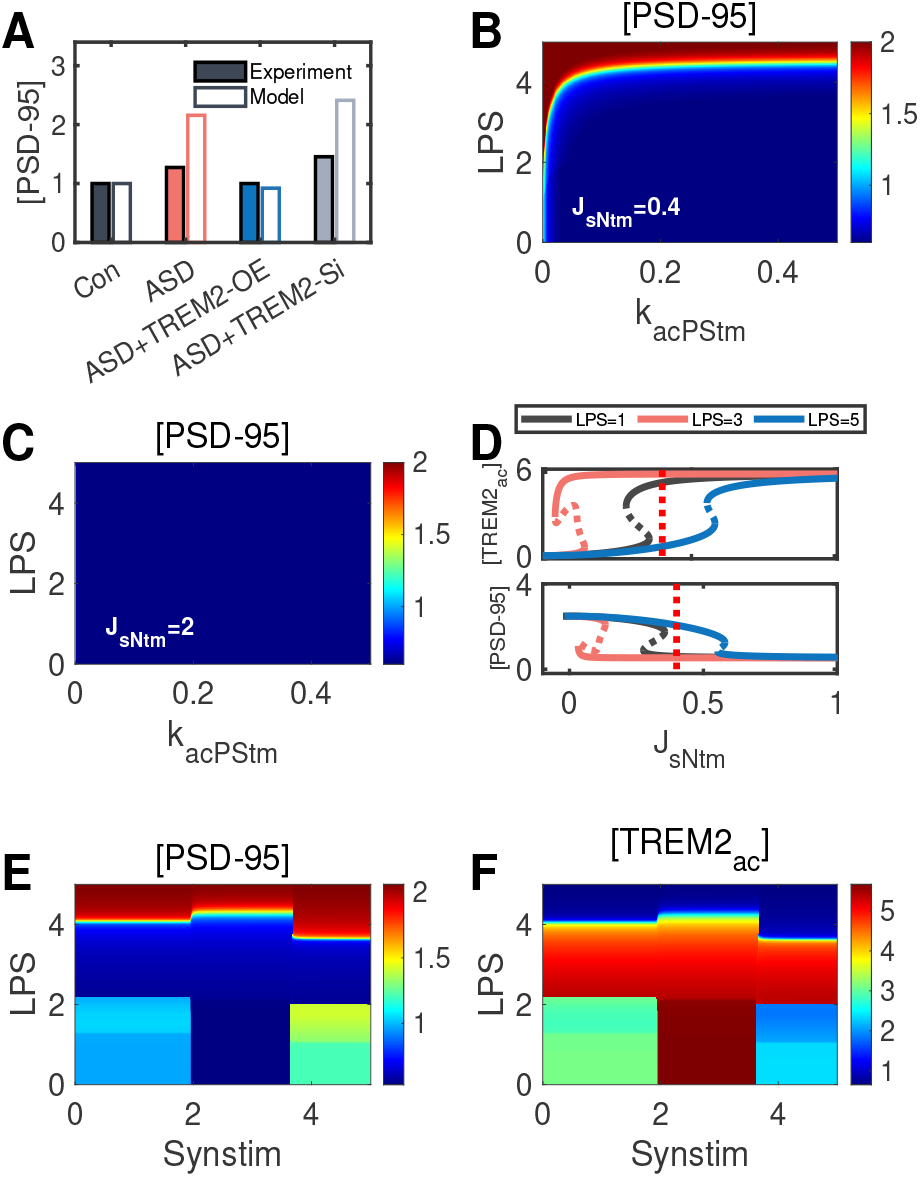
LPS and Synstim co-regulate microglial pruning. **A**. Comparison of experimental and simulated [PSD-95] levels under different conditions: control, ASD, ASD with TREM2 overexpression (ASD+TREM2-OE), and ASD with TREM2 knockdown (ASD+TREM2-si). **B**. Heatmap of [PSD-95] as a function of LPS and *k*_*acPStm*_ at *J*_*sNtm*_ = 0.4. **C**. Heatmap of [PSD-95] as a function of LPS and *k*_*acPStm*_ at *J*_*sNtm*_ = 2. **D**. Bifurcation diagram of [TNFα] and [PSD-95] with respect to *J*_*sNtm*_ under Synstim = 1, 3, and 5. **E**. Heatmap of [PSD-95] as a function of LPS and Synstim. **F**. Heatmap of [TREM2_*ac*_] as a function of LPS and Synstim.

As shown in Figure 7A, [PSD-95] levels in the ASD group were significantly elevated relative to the control group, consistent with both experimental findings^31^ and pathological observations. In the ASD model, overexpression of [TREM2_*tot*_] led to a marked reduction in [PSD-95], approaching normal levels. Conversely, silencing TREM2 resulted in a further elevation of [PSD-95], underscoring the pivotal role of TREM2 in regulating synaptic pruning.

Because the inconsistent feed-forward loops of NFκB-Caspase-3-TREM2_*ac*_and NFκB-TREM2_*tot*_-TREM2_*ac*_would have different effects on pruning of NFκB, We explored the effect of these loops on synaptic pruning by varying *k*_*acPStm*_ and *J*_*sNtm*_(Fig.7(B-D)). As shown in Figure 7B, the very low activation of NFκB at Synstim = 3 and low levels of LPS results in a weak inhibition of TREM2, whereas *k*_*acPStm*_ has a small effect on TREM2, achieving a delicate balance between the two pathways, Normal levels of synaptic pruning were also maintained. If LPS increases, lower *k*_*acPStm*_ results in reduced pruning and the balance is skewed toward NFκB-TREM2_*tot*_-TREM2_*ac*_. However, TREM2_*tot*_ expression was significantly suppressed at high levels of LPS, and microglial pruning remained low regardless of the size of *k*_*acPStm*_. As shown in Fig. 7C, increasing *J*_*sNtm*_ made it more difficult to inhibit TREM2, and changes in LPS or *k*_*acPStm*_ no longer affected synaptic pruning. These results suggest that increasing *J*_*sNtm*_ helps mitigate LPS-induced synaptic accumulation. Figure 7D further validates this result.

As shown in Fig. 7E, [PSD-95] accumulation exhibits a stronger dependence on LPS levels than on synaptic activity. Specifically, at moderate synaptic stimulation (Synstim), [PSD-95] levels remained consistently low across low to moderate

LPS concentrations and did not change with increasing LPS, indicating reduced sensitivity of synapses to LPS under these conditions. However, the combination of low LPS and high Synstim promoted [PSD-95] accumulation at both low and high synaptic activity levels (corresponding to normal and reduced synaptic states, respectively). These findings suggest that enhanced synaptic activity confers resistance to microglia-mediated pruning. Conversely, under high LPS concentrations, [PSD-95] accumulated regardless of Synstim intensity, indicating an absolute block of synaptic pruning and highlighting the pathological impact of excessive microglial activation in ASD.

To further elucidate how LPS and Synstim regulate [TREM2_*ac*_] levels, we generated a heat map of [TREM2_*ac*_] under varying conditions. Using LPS = 1 and Synstim = 1 as the reference, [PSD-95] levels decreased under moderate LPS and Synstim, potentially due to lower synaptic activity requiring pruning or impaired microglial pruning. However, when Synstim remained moderate and LPS was high (LPS = 5), [PSD-95] was not reduced, suggesting a failure of pruning that may contribute to ASD pathology. In contrast, enhanced synaptic activity was associated with elevated and more stable [PSD-95] levels.

Together, the results in Fig.7E–7F indicate a dynamic interplay among LPS concentration, synaptic stimulation, and microglial activation, which shapes the expression patterns of [TREM2_*ac*_] and [PSD-95]. At Synstim = 3, synapses appear particularly sensitive to moderate LPS, while higher Synstim stabilizes [PSD-95]. Full microglial activation under high LPS levels promotes [PSD-95] accumulation, providing mechanistic insight into E/I imbalance in neurodevelopmental disorders such as ASD.

## 4. Discussion and Conclusion

Previous research has demonstrated that prenatal exposure to LPS can induce microglial pro-inflammatory activation and impair synaptic pruning^33^. Based on these findings, we hypothesize that synaptic pruning deficits in ASD are driven by microglial activation and altered synaptic activity. Our results indicate that: (1) Under the stimulus of high levels of LPS, the pruning capability of microglia is decreased, and the synapses that should be pruned are unable to be pruned in time, resulting in an increase in the number of synapses. (2) High levels of synaptic strength to some extent protect against LPS-induced synaptic apoptosis, which may indirectly lead to inadequate synaptic pruning (Fig.4F).

Specifically, as illustrated in Fig.8, there is a non-linear relationship between the pro-inflammatory activation state of LPS-induced microglia and their synaptic pruning capability. At extremely low activation levels (Intermediate proinflammatory state 1, I1-M0/M1), microglial pruning capability remains comparable to the non-activated state. At moderate activation levels (Intermediate proinflammatory state 2, I2-M0/M1), microglial pruning capability improves, leading to reduced synaptic number (as indicated by decreased PSD-95, see Fig.7E). However, at high activation levels (Fully proinflammatory activated state, M1), pruning capability declines, resulting in increased synaptic accumulation. From the perspective of synaptic stimulus, Synstim-induced LTD and LTP have some priority in regulating synaptic pruning, which can be initiated in a noninflammatory state, and synapses are usually the main target of pruning after LTD occurs under moderate stimulus. When microglia are fully activated, their synaptic pruning capability is defective, and the synapses of LTD are not pruned in time, and ultimately excessive accumulation in the brain leads to the pathogenesis of ASD.

**Figure 8.**
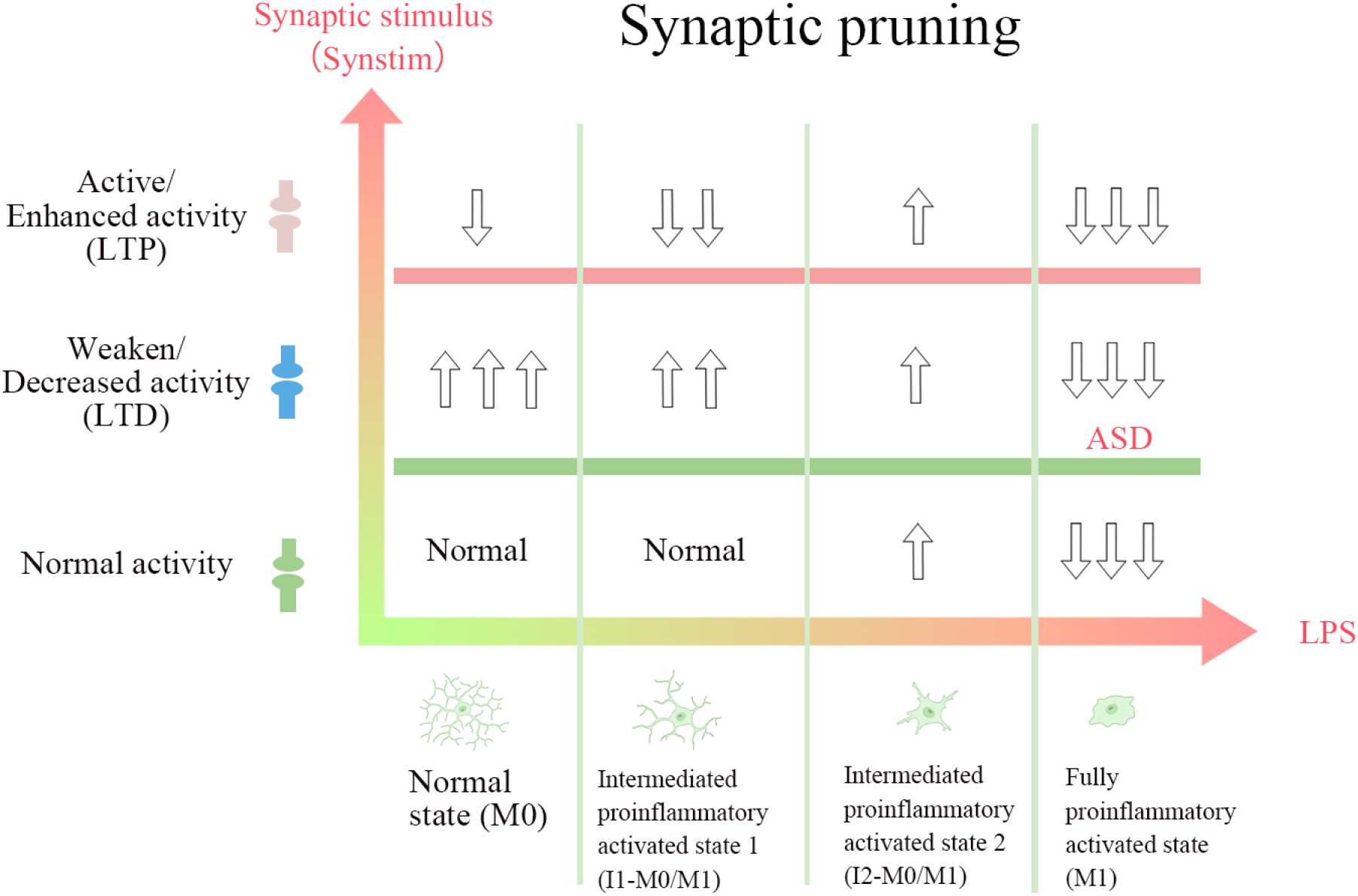
Schematic diagram of changes in the degree of synaptic pruning. The horizontal and vertical axes indicate the levels of change for different concentrations of LPS and different degrees of synaptic stimulus, respectively. When LPS concentrations are highest, synapses that should be pruned fail to be pruned, leading to ASD. Upward arrows indicate an increase relative to normal and downward arrows indicate a decrease relative to normal. The number of arrows indicates the degree of increase or decrease, with more arrows indicating a higher degree.

### Synapse

Typically, microglia sense synaptic activity and modulate the synapse numbers^15^. The changes in synaptic activity are closely associated with the activation of Caspase-3 signaling pathways^7,20^. Downstream effectors of this pathway, such as PtdSer, serve as important tagging for synaptic pruning^7,64,65^. In our work, both CaMKII_*p*_ and PP1 levels can respond to changes in synaptic stimulus, thereby regulating LTD-like synaptic pruning processes^19,74^. Consistent with previous research, our results demonstrate that intense PP1 expression with decreased CaMKII_*p*_ expression, which results in weakened synaptic activity, promotes the occurrence of pruning^19,48,75^. Additionally, once this pathway is blocked, synaptic pruning will not be properly initiated, and then synapses are highly likely to accumulate and cause diseases such as ASD.

From the synaptic perspective, Caspase-3, a key biomarker of synaptic apoptosis, plays a central role in initiating synaptic pruning. XIAP, an upstream inhibitor of Caspase-3, has been shown to suppress Caspase-3 activation; its overexpression can lead to impaired synaptic elimination and promote synaptic accumulation, contributing to the pathophysiology of ASD^76,77^. Therefore, in ASD cases characterized by excessive synapses, targeted modulation of the XIAP–Caspase-3 axis may offer a promising therapeutic strategy. Since both TNFα and weaken synaptic activity facilitate Caspase-3 activation, adjusting synaptic stimulation levels and LPS exposure may provide an effective means to regulate this pathway.

Moreover, it is noteworthy that enhanced synaptic activity can paradoxically exacerbate synaptic accumulation when microglial function is compromised. This observation is consistent with experimental findings indicating that increased excitatory drive is a hallmark of ASD^78,79^.

### Microglia

Previous studies have shown that reduced TREM2 expression strongly correlates with the onset of ASD and the severity of symptoms^26^. TREM2 deficiency leads to decreased microglia phagocytosis, and its upregulation inhibits pro-inflammatory responses^31,38^. Our model suggests that the degree of pro-inflammatory microglia activation can largely influence the synaptic pruning capability and the degree of pruning. To be specific, fully proinflammatory activated microglia are not able to prune the weakened synapse for its decreased pruning capability, which directly results in the accumulation of synapses that need to be pruned that cannot be eliminated by pruning, which may be related to the increase in immature spines in pathological studies^26^.

### Interventions and treatments

The interaction of microglia with neurons determines which synapses are pruned. Thus, we think that the interaction process between the two needs to be considered in the implementation of intervention and treatment. Our study suggests that when synaptic pruning should be initiated, the pruned synapses are not pruned because of the abnormal pruning capability of microglia, which further corroborates the idea that the interaction between microglia and neurons determines whether synapses are pruned or not. Our results provide new ideas for the intervention and treatment of ASD, that is, by directly or indirectly regulating the pruning capability of microglia, or by regulating synaptic strength (synaptic excitability).

Combined with our model, we think that interventions and treatments of ASD can be implemented from four perspectives to address synaptic abnormalities in ASD: (1) By using anti-inflammatory drugs, microglia can be changed from a fully proinflammatory activated state to a mildly activated state or even to a normal state, so that TREM2 can be changed from a low state to a high state (Fig.2C), so as to enhance its pruning capability and reduce the accumulation of synapses. Alternatively, the activator of TREM2 was added directly to enhance the synaptic pruning capability of microglia. (2) In the case of quiescent or incompletely pro-inflammatory activated microglia (I2-M0/M1), the synaptic activity is appropriately reduced through chemical or electrical stimulation, thereby enhancing the synaptic pruning level of microglia. (3) Appropriate and localized activation of Caspase-3, for example through optical stimulation, thereby facilitating synapse-intrinsic apoptotic signaling and reducing synaptic overaccumulation. (4) By applying inhibitory neurotransmitters and reducing the excitability of brain circuits related to ASD, it may help alleviate the symptoms of ASD.

## Limitations

In addition to the TREM2–PtdSer pathway explored in this study, other signaling pathways such as CX3CL1– CX3CR1, C1q/C3, and CD47–SIRPα have also been implicated in regulating synaptic pruning. These pathways primarily function as interfaces for communication between synapses and microglia, helping establish recognition and contact. However, the downstream molecular events within microglia or synapses triggered by these pathways remain largely uncharacterized in experimental studies. For example, although the binding of synaptic CX3CL1 to microglial CX3CR1 is known to influence pruning, it is still unclear what intracellular signaling cascades are subsequently activated to drive engulfment. Similarly, the role and mechanisms of the CD47–SIRPα pathway in ASD are poorly understood.

Given this limited mechanistic clarity and to maintain tractability in our modeling, we did not incorporate these pathways. Additionally, while astrocytes have been shown to participate in synaptic engulfment^80^, they were not considered in our framework. This is because microglia are recognized to carry out synaptic pruning with greater precision and efficiency. When this pruning function becomes dysregulated under inflammatory conditions, it can lead to pronounced neural circuit abnormalities^81^. Therefore, our results provide a simplified yet informative perspective, offering a basis for future exploration into the complex, multi-pathway regulation of synaptic pruning.

In summary, our model proposes a potential pathogenesis of ASD from the perspective of synaptic pruning: synapses that should be pruned accumulate due to the reduced pruning capability of microglia, leading to an increased number of synapses and, consequently, the development of ASD. We also emphasize the critical role of microglial pro-inflammatory activation in the pruning process and the guiding effect of synaptic activity on synaptic pruning. Our work, while qualitatively consistent with existing experimental findings, offers additional predictions that may provide insights into the treatment of ASD.

## Acknowledgements (not compulsory)

Acknowledgements should be brief, and should not include thanks to anonymous referees and editors, or effusive comments. Grant or contribution numbers may be acknowledged.

## Author contributions statement

Must include all authors, identified by initials, for example: A.A. conceived the experiment(s), A.A. and B.A. conducted the experiment(s), C.A. and D.A. analysed the results. All authors reviewed the manuscript.

