## Supporting information for "Dysfunction of microglia-mediated synaptic pruning in Autism spectrum disorder"

### Supplementary information

July 21, 2025

#### 1 Equations

$$[NF\kappa B] = [NF\kappa B_{\mathrm{tot}}] - [NF\kappa B_{\mathrm{p}}] \tag{1}$$

$$\begin{aligned}\frac{d[NF\kappa B_{\mathrm{p}}]}{dt} &= k_{acLPSn} \cdot LPS \cdot \frac{[NF\kappa B]}{[NF\kappa B] + J_{acLPSn}} \\ &+ k_{acTNFn} \cdot \frac{[TNF\alpha]^4}{[TNF\alpha]^4 + J_{an}^4} \cdot \frac{[NF\kappa B]}{[NF\kappa B] + J_{acTNFn}} \\ &- k_{dpn} \cdot \frac{[NF\kappa B_{\mathrm{p}}]}{[NF\kappa B_{\mathrm{p}}] + J_{dpn}} - k_{dpTREM2n} \cdot [TREM2_{ac}] \cdot \frac{[NF\kappa B_{\mathrm{p}}]}{[NF\kappa B_{\mathrm{p}}] + J_{dpTREM2n}}\end{aligned} \tag{2}$$

$$\frac{d[TNF\alpha]}{dt} = k_{s0ta} + k_{sNta} \cdot \frac{[NF\kappa B_p]^2}{[NF\kappa B_p]^2 + J_{sNta}^2} - k_{dta} \cdot [TNF\alpha] \tag{3}$$

$$\begin{aligned}\frac{d[TREM2_{ac}]}{dt} &= k_{0actm} + k_{acPStm} \cdot \frac{[ePtdSer]^2}{[ePtdSer]^2 + J_{acPStm}^2} \cdot ([TREM2_{tot}] - [TREM2_{ac}]) \\ &- k_{dtmac} \cdot [TREM2_{ac}]\end{aligned} \tag{4}$$

$$\frac{d[TREM2_{tot}]}{dt} = k_{s0tm} + k_{sNtm} \cdot \frac{J_{sNtm}^2}{[NF\kappa B_{\mathrm{p}}]^2 + J_{sNtm}^2} - k_{dtmtot} \cdot [TREM2_{tot}] \tag{5}$$

$$\begin{aligned}\frac{d[XIAP]}{dt} &= k_{s0iap} + k_{sCMKiap} \cdot \frac{[CaMKII_{\mathrm{p}}]}{[CaMKII_{\mathrm{p}}] + J_{sCMKiap}} \\ &- k_{dBaxiap} \cdot \frac{[Bax]^4}{[Bax]^4 + J_{xx}^4} \cdot \frac{[XIAP]}{[XIAP] + J_{Baxiap}} - k_{diap} \cdot [XIAP]\end{aligned} \tag{6}$$

$$\begin{aligned}\frac{d[Bax]}{dt} &= \left(k_{0acx} + k_{acTNFx} \cdot \frac{[TNF\alpha]^4}{[TNF\alpha]^4 + {J_{acTNFx}}^4} + k_{acPP1x} \cdot \frac{[PP1]^4}{[PP1]^4 + {J_{acPP1x}}^4}\right) \\ &\cdot ([Bax_{tot}] - [Bax]) - k_{dx} \cdot [Bax]\end{aligned} \tag{7}$$

$$\begin{aligned}\frac{d[CytoC]}{dt} &= \left(k_{0accc} + k_{accc} \cdot [Bax] \cdot \frac{[Casp3]^4}{[Casp3]^4 + J_{accc}^4}\right) \cdot ([CytoC_{tot}] - [CytoC]) \\ &- k_{dcc} \cdot [CytoC]\end{aligned} \tag{8}$$

$$\begin{aligned}\frac{d[Casp3]}{dt} &= \left(k_{0acc3} + k_{acc3} \cdot \frac{[CytoC]^4}{[CytoC]^4 + J_{acc3}^4}\right) \cdot ([Casp3_{tot}] - [Casp3]) \\ &- k_{0dc3} \cdot [Casp3] - k_{diapc3} \cdot [XIAP] \cdot [Casp3]\end{aligned} \tag{9}$$

$$\frac{d[CaCaM]}{dt} = k_{0sca} + k_{sSca} \cdot \frac{Synstim^4}{Synstim^4 + J_{sSca}^4} - k_{dca} \cdot [CaCaM] \tag{10}$$

$$\frac{d[CaMKII_p]}{dt} = k_{acCack} \cdot \frac{[CaCaM]^4}{[CaCaM]^4 + J_{acCack}^4} \cdot \frac{[CaMKII]}{[CaMKII] + J_{ck}} + k_{acCMKck} \cdot [CaMKII_p] \cdot \frac{[CaMKII]}{[CaMKII] + J_{acCMKck}} + k_{ac0ck} \cdot [CaMKII_{p0}] - k_{dpck} \cdot ([PP1] + [PP1_0]) \cdot \frac{[CaMKII_p]}{[CaMKII_p] + J_{dpck}} \quad (11)$$

$$[CaMKII] = [CaMKII_{tot}] - [CaMKII_p] \quad (12)$$

$$\frac{d[PP1]}{dt} = k_{dCapp} \cdot \frac{[CaCaM]^8}{[CaCaM]^8 + J_{dCapp}^8} \cdot \frac{[PP1_p]}{[PP1_p] + J_{pp}} + k_{dP1pp} \cdot [PP1] \cdot \frac{[PP1_p]}{[PP1_p] + J_{dP1pp}} + k_{d0pp} \cdot [PP1_0] - k_{acCMKpp} \cdot ([CaMKII_p] + [CaMKII_{p0}]) \cdot \frac{[PP1]}{[PP1] + J_{acCMKpp}} \quad (13)$$

$$[PP1_p] = [PP1_{tot}] - [PP1] \quad (14)$$

$$\frac{d[ePtdSer]}{dt} = k_{acC3ps} \cdot [PtdSer] \cdot \frac{[Casp3]^2}{[Casp3]^2 + J_{acC3ps}^2} - k_{dps} \cdot [ePtdSer] \quad (15)$$

$$\frac{d[PSD95]}{dt} = k_{s05} - k_{dtTREM25} \cdot [TREM2_{ac}] \cdot \frac{[PSD95]}{[PSD95] + J_{dtTREM25}} - k_{d5} \cdot [PSD95] \quad (16)$$

#### 2 Supplemental Tables

**Table 1: Variables and their initial values**

| Variable | Description | Initial values |
|---|---|---|
| $[TNF\alpha]$ | Concentration of $TNF\alpha$ | 0.017 |
| $[TREM2_{ac}]$ | Concentration of active TREM2 | 3.33 |
| $[TREM2_{tot]}$ | Concentration of total TREM2 | 6.02 |
| $[XIAP]$ | Concentration of XIAP | 0.67 |
| $[ePtdSer]$ | Concentration of ePtdSer | 0.163 |
| $[Bax]$ | Concentration of Bax | 0.00033 |
| $[CytoC]$ | Concentration of CytoC | 0.0589 |
| $[Casp3]$ | Concentration of Casp3 | 0.15 |
| $[PSD95]$ | Concentration of PSD95 | 0.99 |
| $[CaM]$ | Concentration of CaM | 0.017 |
| $[CaMKII_p]$ | Concentration of $\text{CaMKII}_p$ | 0.0411 |
| $[PP1]$ | Concentration of PP1 | 0.00165 |

**Table 2: Parameters for the whole model**

| Parameter | Description | Value | References |
|---|---|---|---|
| $k_{s0ta}$ | Basal production rate of $TNF\alpha$ | 0.0001 | [1] |
| $k_{sNta}$ | $NF\kappa$B-dependent production rate of $TNF\alpha$ | 0.02 | [1] |
| $J_{sNta}$ | Michaelis constant of $NF\kappa$B-dependent production | 0.2 | Assumed |
| $k_{dta}$ | Basal degradation rate of $TNF\alpha$ | 0.006 | Assumed |
| NF$\kappa$B$_{tot}$ | The total concentrations of all forms of NF$\kappa$B | 1.5 | Assumed |
| $k_{acLPSn}$ | LPS-dependent phosphorylation rate of $NF\kappa$B | 0.04 | [2] |
| $J_{acLPSn}$ | Michaelis constant of LPS-dependent $NF\kappa$B phosphorylation | 1 | Assumed |
| $k_{acTNFn}$ | $TNF\alpha$-dependent phosphorylation rate of $NF\kappa$B | 0.06 | [3, 4] |
| $J_{acTNFn}$ | Michaelis constant of $TNF\alpha$-dependent $NF\kappa$B phosphorylation | 0.8 | Assumed |
| $J_{an}$ | Michaelis constant for the binding of $TNF\alpha$ to its ligand on microglia | 0.3 | Assumed |
| $k_{dpn}$ | Basal dephosphorylation rate of $NF\kappa$B | 0.06 | Assumed |
| $J_{dpn}$ | Michaelis constant of basal $NF\kappa$B phosphorylation | 0.2 | Assumed |
| $k_{dpTREM2n}$ | $TREM2_{ac}$-dependent dephosphorylation rate of $NF\kappa$B | 0.04 | [5] |
| $J_{dpTREM2n}$ | Michaelis constant of $TREM2_{ac}$-dependent $NF\kappa$B dephosphorylation | 0.2 | Assumed |
| $k_{0actm}$ | Basal activation rate of $TREM2_{ac}$ | 0.002 | Assumed |
| $k_{acPStm}$ | $PtdSer$-dependent activation rate of $TREM2_{ac}$ | 0.1 | [6, 7] |
| $J_{acPStm}$ | Michaelis constant of $PtdSer$-dependent $TREM2_{ac}$ activation | 0.6 | Assumed |
| $k_{dtmac}$ | Basal deactivation rate of $TREM2_{ac}$ | 0.006 | Assumed |
| $k_{0stm}$ | Basal induction rate of $TREM2_{tot}$ | 0.0001 | Assumed |
| $k_{sNtm}$ | $NF\kappa$B-dependent inhibition rate of $TREM2_{tot}$ production | 0.03 | [8] |
| $J_{sNtm}$ | Michaelis constant of $NF\kappa$B-dependent inhibition of $TREM2_{tot}$ production | 0.4 | Assumed |
| $k_{dtmtot}$ | Basal degradation rate of $TREM2_{tot}$ | 0.005 | Assumed |
| $k_{s0iap}$ | Basal induction rate of $XIAP$ | 0.1 | Assumed |
| $k_{sCMKiap}$ | $CaMKII_p$-dependent production rate of $XIAP$ | 0.06 | Assumed |
| $J_{sCMKiap}$ | Michaelis constant of $CaMKII_p$-dependent production of $XIAP$ | 2 | Assumed |
| $k_{diap}$ | Basal degradation rate of $XIAP$ | 0.15 | Assumed |
| $k_{dBaxiap}$ | Bax-dependent inhibition rate of $XIAP$ | 0.07 | Assumed |

**Table 2: Parameters for the whole model**

| Parameter | Description | Value | References |
|---|---|---|---|
| $J_{xx}$ | Michaelis constant of Bax-dependent inhibition of $XIAP$ | 1 | Assumed |
| $J_{dBaxiap}$ | Michaelis constant of Bax-dependent degradation of $XIAP$ | 0.4 | Assumed |
| $k_{0acx}$ | Basal activation rate of $Bax$ | 0.0001 | Assumed |
| $k_{acTNFx}$ | $TNF\alpha$-dependent activation rate of $Bax$ | 0.3 | [9] |
| $J_{acTNFx}$ | Michaelis constant of $TNF\alpha$-dependent $Bax$ activation | 3 | Assumed |
| $k_{acPP1x}$ | $PP1$-dependent activation rate of $Bax$ | 0.2 | [10, 11, 12] |
| $J_{acPP1x}$ | Michaelis constant of $PP1$-dependent $Bax$ activation | 3 | Assumed |
| $k_{dx}$ | Basal deactivation rate of $Bax$ | 0.3 | Assumed |
| $Bax_{tot}$ | The total concentrations of all forms of Bax | 1 | Assumed |
| $k_{0accc}$ | Basal release rate of $CytoC$ | 0.001 | Assumed |
| $k_{accc}$ | $Bax$-dependent release rate of $CytoC$ | 0.9 | [13] |
| $J_{accc}$ | Michaelis constant of $Bax$-dependent $CytoC$ release | 0.9 | Assumed |
| $k_{dcc}$ | Basal degradation rate of $CytoC$ | 0.05 | Assumed |
| $CytoC_{tot}$ | The total concentrations of all forms of $CytoC$ | 3 | Assumed |
| $k_{0acc3}$ | Basal activation rate of $Casp3$ | 0.001 | Assumed |
| $k_{acc3}$ | $CytoC$-dependent activation rate of $Casp3$ | 0.9 | [13] |
| $J_{acc3}$ | Michaelis constant of $CytoC$-dependent $Casp3$ activation | 0.3 | [13] |
| $k_{0dc3}$ | Basal deactivation rate of $Casp3$ | 0.01 | Assumed |
| $k_{diapc3}$ | $XIAP$-dependent degradation rate of $Casp3$ | 0.05 | [14] |
| $Caspase3_{tot}$ | The total concentrations of all forms of $Caspase3$ | 3 | Assumed |
| $k_{0acps}$ | Basal externalisation rate of $PtdSer$ | 0.001 | Assumed |
| $k_{acC3ps}$ | $Casp3$-dependent externalisation rate of $PtdSer$ | 0.6 | Assumed |
| $J_{acC3ps}$ | Michaelis constant of $Casp3$-dependent $PtdSer$ externalisation | 2 | Assumed |
| $k_{dps}$ | Basal internalisation rate of $PtdSer$ | 0.1 | Assumed |
| $PtdSer_{tot}$ | The total concentrations of all forms of $PtdSer$ | 3 | Assumed |
| $k_{s05}$ | Basal production rate of PSD-95 | 0.1 | Assumed |
| $k_{dtTREM25}$ | $TREM2_{ac}$-dependent degradation rate of PSD-95 | 0.04 | [15] |
| $J_{dtTREM25}$ | Michaelis constant of $TREM2_{ac}$-dependent PSD-95 degradation | 1 | Assumed |
| $k_{d5}$ | Basal degradation rate of PSD-95 | 0.04 | Assumed |
| $k_{acCack}$ | $CaCaM$-dependent phosphorylation rate of $CaMKII$ | 120 | [16] |
| $J_{acCack}$ | Michaelis constant of combination of $CaMKII$ and $CaCaM$ | 4 | [16] |

**Table 2: Parameters for the whole model**

| Parameter | Description | Value | References |
|---|---|---|---|
| $J_{ck}$ | Michaelis constant of $CaCaM$-dependent phosphorylation | 0.1 | Assumed |
| $k_{acCMKck}$ | Autophosphorylation rate of $CaMKII$ | 2 | [16] |
| $J_{acCMKck}$ | Michaelis constant of autophosphorylation of $CaMKII$ | 15 | Assumed |
| $k_{ac0ck}$ | Basal activated rate of $CaMKII_p$ | 1 | Assumed |
| $k_{dpck}$ | $PP1$-dependent dephosphorylation rate of $CaMKII_p$ | 30 | [16] |
| $J_{dpck}$ | Michaelis constant of $PP1$-dependent dephosphorylation of $CaMKII_p$ | 1 | Assumed |
| $CaMKII_{tot}$ | The total concentrations of all forms of $CaMKII_p$ | 5 | Assumed |
| $k_{dCapp}$ | $CaCaM$-dependent phosphorylation rate of $PP1_p$ | 80 | [16] |
| $J_{dCapp}$ | Michaelis constant of combination of $PP1_p$ and $CaCaM$ | 0.8 | Assumed |
| $J_{pp}$ | Michaelis constant of $CaCaM$-dependent $PP1_p$ phosphorylation | 3 | Assumed |
| $k_{dP1pp}$ | Autodephosphorylation rate of $PP1_p$ | 2 | [16] |
| $J_{dP1pp}$ | Michaelis constant of autodephosphorylation of $PP1_p$ | 5 | Assumed |
| $k_{d0pp}$ | Basal activated rate of $PP1$ | 1 | Assumed |
| $k_{acCMKpp}$ | $CaMKII_p$-dependent phosphorylation rate of $PP1$ | 15 | Assumed |
| $J_{acCMKpp}$ | Michaelis constant of $CaMKII_p$-dependent phosphorylation of $PP1$ | 0.01 | Assumed |
| $\text{PP1}_{tot}$ | The total concentrations of all forms of PP1 | 3 | Assumed |
| $k_{0sca}$ | Basal $Ca^{2+}$ influx production rate | 0.001 | Assumed |
| $k_{sSca}$ | $S$-dependent production rate of $Ca^{2+}$ influx | 2 | Assumed |
| $J_{sSca}$ | Michaelis constant of $S$-dependent production of $Ca^{2+}$ influx | 5 | Assumed |
| $k_{dca}$ | Basal $Ca^{2+}$ efflux rate | 0.1 | Assumed |
| $\text{NMDAR}_{tot}$ | The total concentrations of all forms of NMDAR | 1 | Assumed |

[8] Rosie Owens, Kathleen Grabert, Claire L. Davies, Alessio Alfieri, Jack P. Antel, Luke M. Healy, and Barry W. McColl. Divergent neuroinflammatory regulation of microglial trem expression and involvement of nf- b. *Frontiers in Cellular Neuroscience*, 11, March 2017.

#### 3 Supplemental Figures: Sensitivity analysis

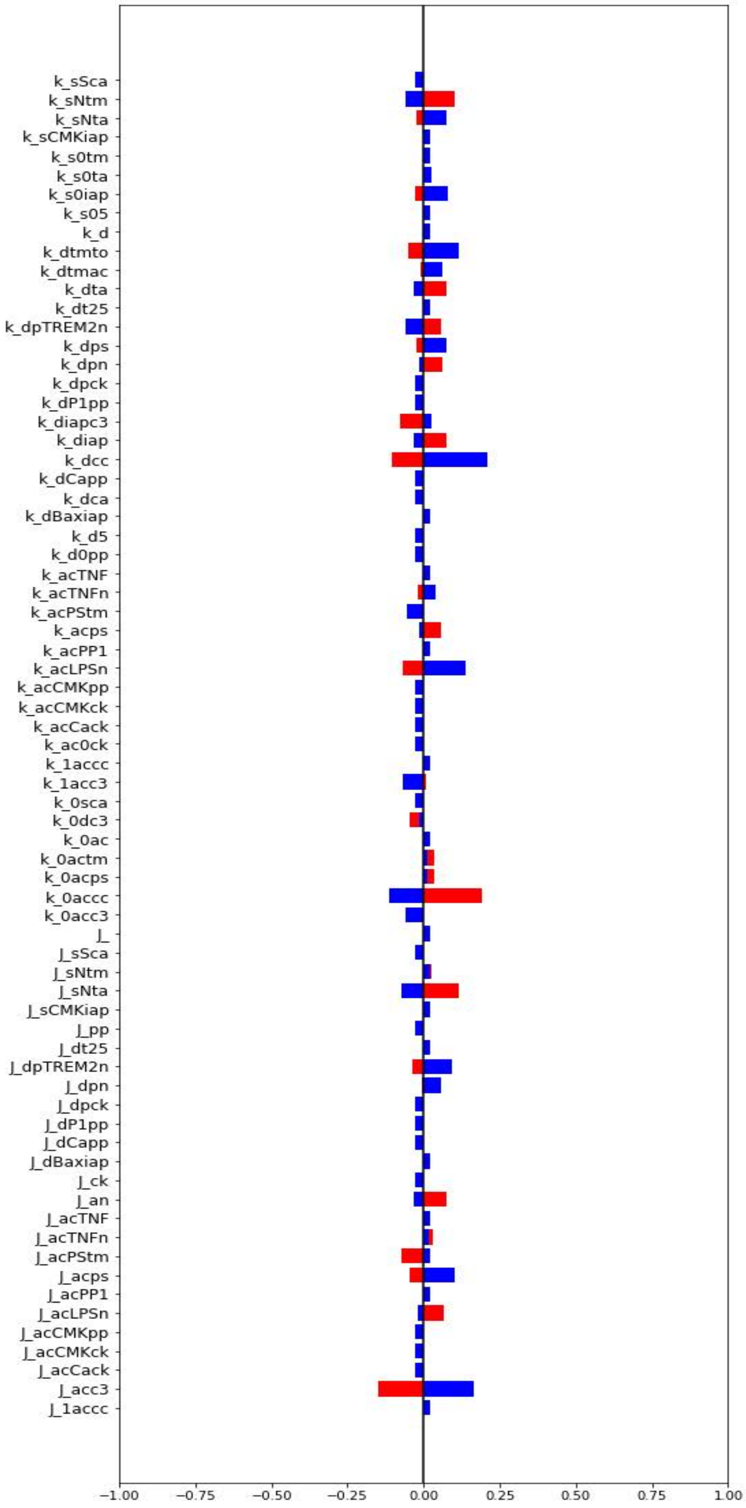


Figure 1: Sensitivity analysis of the bifurcation point SN1

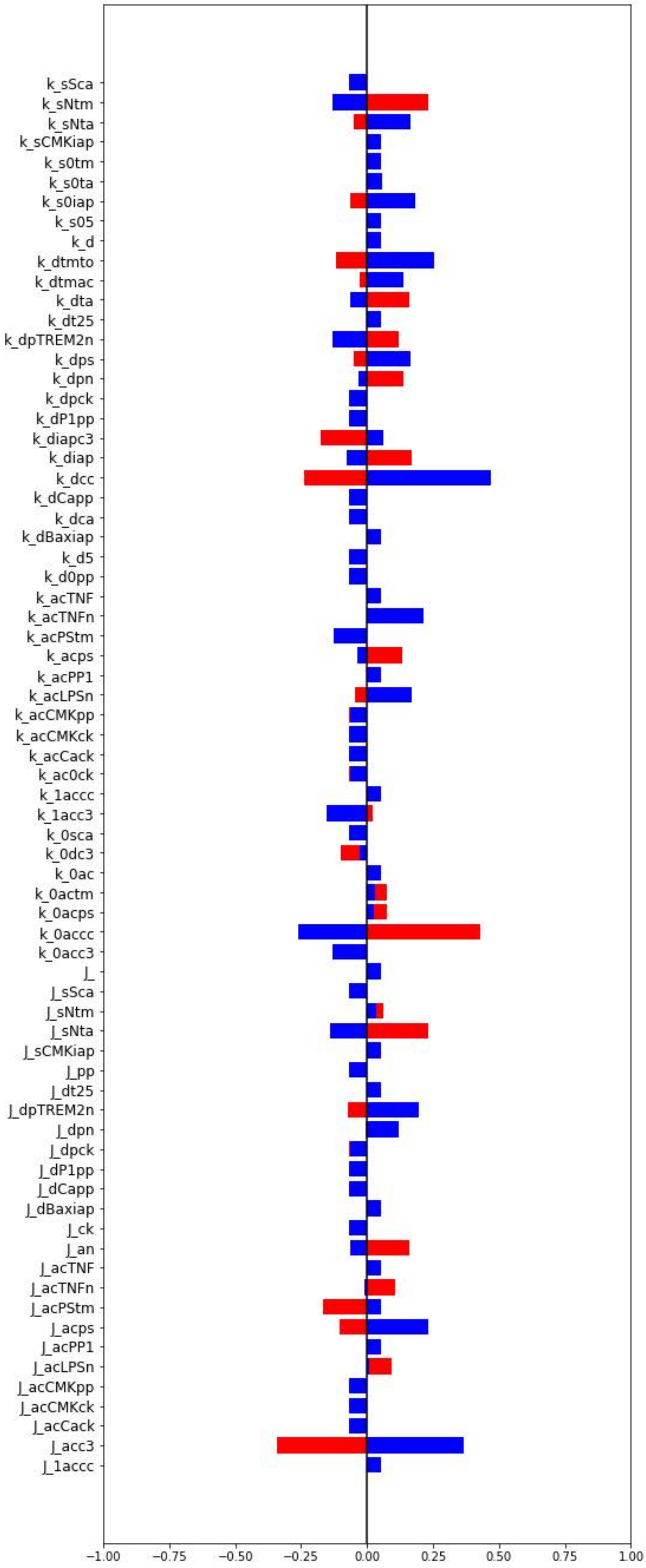


Figure 2: Sensitivity analysis of the bifurcation point SN2

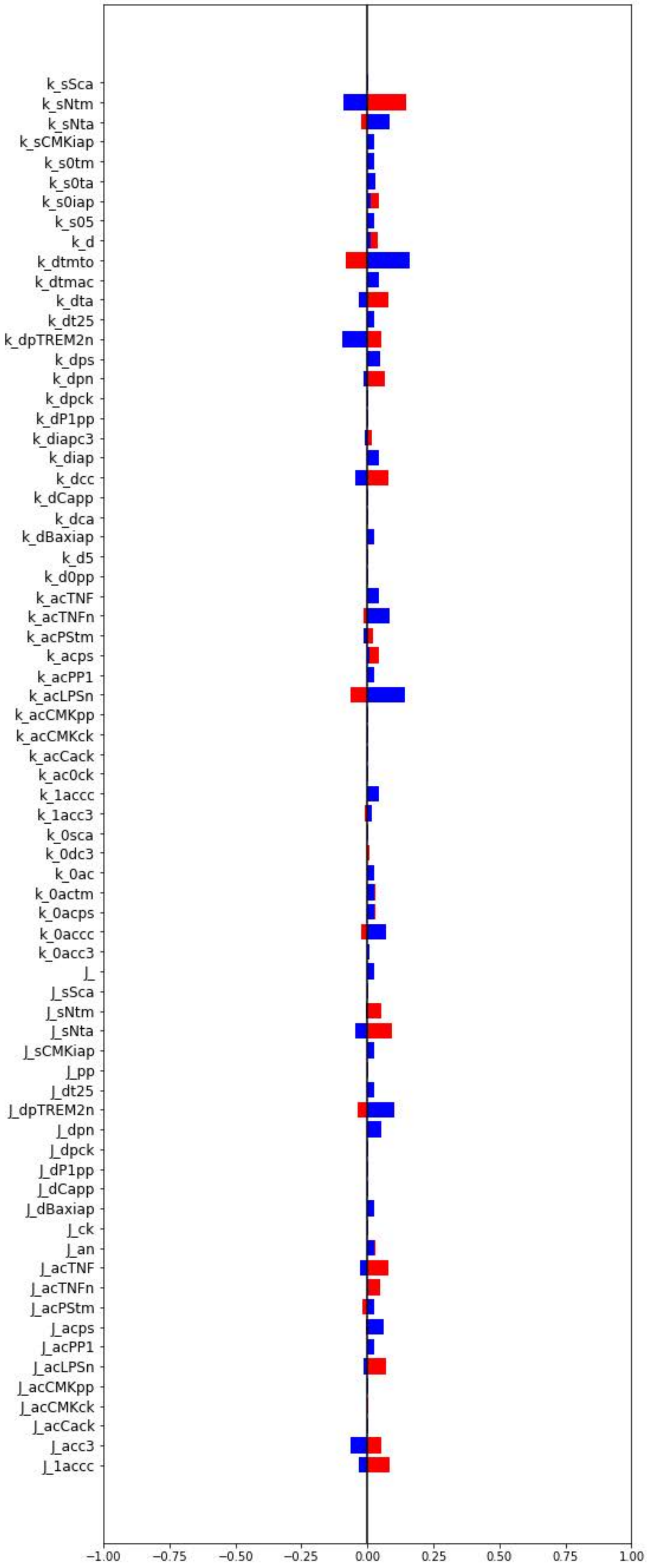


Figure 3: Sensitivity analysis of the bifurcation point SN3

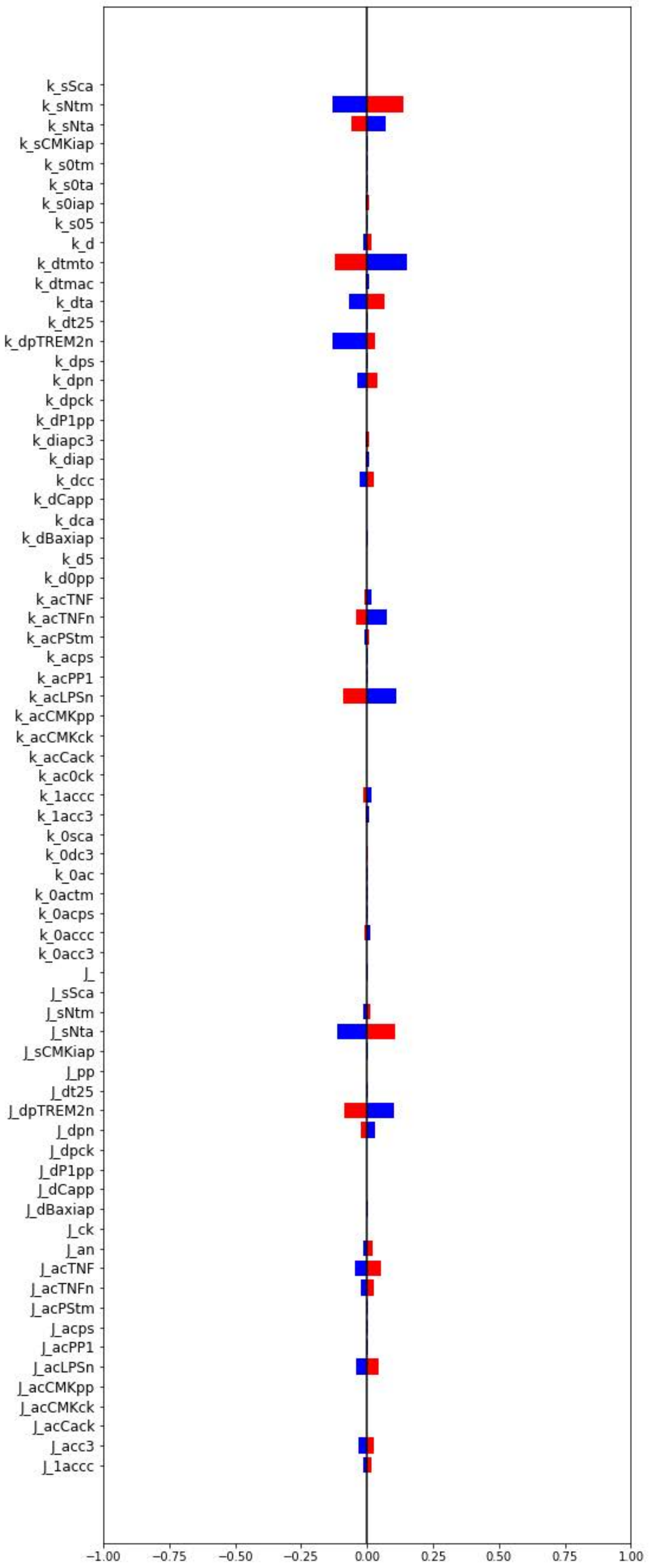


Figure 4: Sensitivity analysis of the bifurcation point SN4

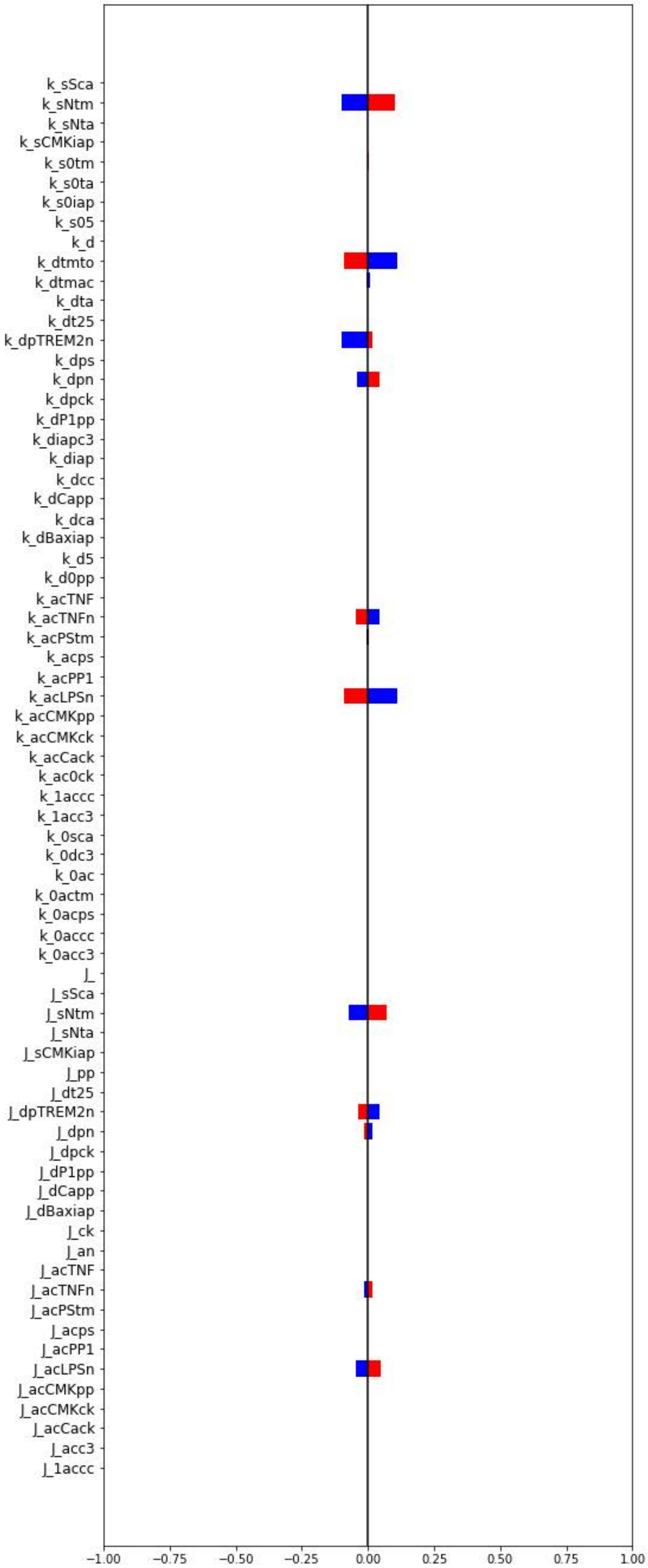


Figure 5: Sensitivity analysis of the bifurcation point SN5

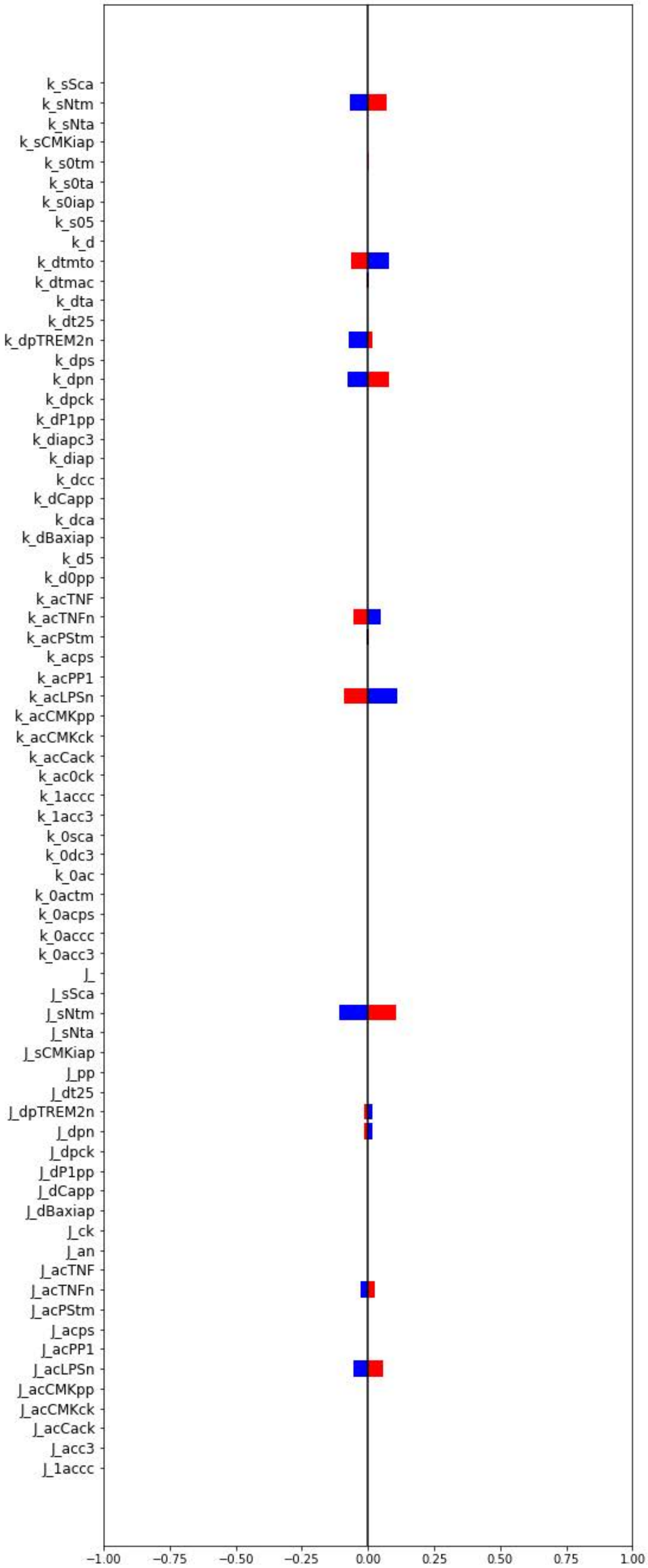


Figure 6: Sensitivity analysis of the bifurcation point SN6
